# TIR domains of plant immune receptors are 2′,3′-cAMP/cGMP synthetases mediating cell death

**DOI:** 10.1101/2021.11.09.467869

**Authors:** Dongli Yu, Wen Song, Eddie Yong Jun Tan, Li Liu, Yu Cao, Jan Jirschitzka, Ertong Li, Elke Logemann, Chenrui Xu, Shijia Huang, Aolin Jia, Xiaoyu Chang, Zhifu Han, Bin Wu, Paul Schulze-Lefert, Jijie Chai

## Abstract

2′,3′-cAMP is a positional isomer of the well-established second messenger 3′,5′-cAMP, but little is known on the biology of this noncanonical cyclic nucleotide monophosphate (cNMP). Toll/interleukin-1 receptor (TIR) domains of nucleotide-binding leucine-rich repeat (NLR) immune receptors have NADase function necessary but insufficient to activate plant immune responses. Here we show that plant TIR proteins, besides being NADases, act as 2′,3′-cAMP/cGMP synthetases by hydrolyzing RNA/DNA. Structural data shows that a TIR domain adopts distinct oligomers with dual and exclusive enzymatic activity. Mutations specifically disrupting the synthetase activity abrogate TIR-mediated cell death in *Nicotiana benthamiana*, supporting an important role for these cNMPs in TIR signaling. Furthermore, the *Arabidopsis* negative regulator of TIR-NLR signaling, NUDT7 displays 2′,3′-cAMP/cGMP but not 3′,5′-cAMP/cGMP phosphodiesterase activity and suppresses cell death activity of TIRs in *N. benthamiana*. Our study identifies a novel family of 2′,3′-cAMP/cGMP synthetase and establishes a role for the noncanonical cNMPs in plant immune responses.

## INTRODUCTION

2′,3′-cAMP is a regioisomer of the canonical intracellular second messenger 3′,5′-cAMP. In contrast to the canonical cAMP, however, little is known about 2′,3′-cAMP. It was not until 2009 and 2014 that the noncanonical cyclic nucleotide monophosphate (cNMP) was identified in animals (Ren et al., 2009) and in plants (Van Damme et al., 2014), respectively. Occurrence of 2′,3′-cAMP and other 2′,3′-cNMPs (2′,3′-cGMP, 2′,3′-cCMP, and 2′,3′-cUMP) has now been demonstrated in bacteria, plants, mice and humans (Jackson, 2017). mRNA turnover by RNases can result in the generation of 2′,3′-cNMPs (Jackson, 2017). Studies in animals support a physiological role for 2′,3′-cNMPs in the response to injury (Azarashvili et al., 2009; Jackson et al., 2014). In *Arabidopsis*, wounding (Van Damme et al., 2014), heat and dark stress conditions (Kosmacz et al., 2018) induce the accumulation of cellular 2′,3′-cAMP/cGMP, demonstrating a correlation of increased 2′,3′-cNMP levels with plant responses to abiotic stress. This notion is strengthened by the observations that 2′,3′-cAMP mediates stress granule (SG) formation (Kosmacz et al., 2018) and mimics the abiotic stress response in *Arabidopsis* (Chodasiewicz et al., 2021). Metabolism of 2′,3′-cNMPs to 2′-NMPs, 3′-NMPs has been demonstrated in animals (Trapp et al., 1988) and plants (Genschik et al., 1997; Tyc et al., 1987), suggesting negative regulations of the noncanonical cNMPs.

Plant defense against microbial pathogens is built on a two-tiered immune system that consists of interdependent pathogen-associated molecular pattern (PAMP)-triggered immunity (PTI) and effector-triggered immunity (ETI) (Ngou et al., 2021; Yuan et al., 2021). ETI is typically mediated by intracellular immune receptors, called nucleotide binding (NB) leucine-rich repeat receptors (NLRs), which recognize strain-specific pathogen effector molecules delivered inside plant cells. NLRs can be largely divided into two groups, with an N-terminal coiled-coil (CC) domain or an N-terminal Toll/interleukin 1 receptor (TIR) domain. Recognition of pathogen effectors results in the formation of oligomeric NLR complexes termed resistosomes (Xiong et al., 2020), as demonstrated for the CC-NLR (CNL) ZAR1 (Wang et al., 2019a) and the TIR-NLRs (TNLs) RPP1 (Ma et al., 2020) and Roq1 (Martin et al., 2020). Activation of NLR resistosomes induces ETI responses, often including a hypersensitive response (HR), a form of regulated cell death that is localized to sites of attempted pathogen infection. Whereas the ZAR1 resistosome is a Ca^2+^-permeable cation channel (Bi et al., 2021), the RPP1 and Roq1 resistosomes function as holoenzymes of NADase encoded within the TIR domain (Horsefield et al., 2019; Ma et al., 2020; Martin et al., 2020; Wan et al., 2019). The TIR-encoded NADase activity is required for activation of immune signaling nodes, EDS1-SAG101 and EDS1-PAD4 heterodimers, and the ‘helper’ NLRs (RNLs) ADR1s and NRG1s (Castel et al., 2019; Lapin et al., 2019; Peart et al., 2005; Qi et al., 2018; Wu et al., 2019). Once TNLs are activated, EDS1-PAD4 and EDS1-SAG101 form hetero-complexes with ADR1s and NRG1s, respectively (Sun et al., 2021; Wu et al., 2021), inducing Ca^2+^-channel activity of the RNLs (Jacob et al., 2021). ADR1 is also important for defense responses of some CNLs, such as *Arabidopsis* RPS2 (Bhandari et al., 2019; Bonardi et al., 2011; Castel et al., 2019; Venugopal et al., 2009; Wu et al., 2019), suggesting cross-talk between CNL and TNL signaling pathways. In addition to immunity, EDS1 signaling is also involved in abiotic stress responses of plants (Suzuki et al., 2012; Wiermer et al., 2005). In *Arabidopsis*, EDS1 signaling is negatively regulated by *AtNUDT7*, a nucleoside diphosphate-linked moiety X (Nudix) hydrolase, and its closest homolog NUDT6 (Bartsch et al., 2006; Ge et al., 2007), although the underlying mechanism, including the associated hydrolase substrates, remains elusive.

In addition to canonical TNLs (TIR-NB-LRR), plant genomes encode many TIR proteins consisting only of a TIR domain (TIR-only or TX) or TIR-NB (TN) with some of them known to have a function in plant defense (Collier et al., 2011; Gao et al., 2018; Meyers et al., 2002; Nandety et al., 2013; Nishimura et al., 2017). For example, the TIR-only protein RBA1 in *Arabidopsis* Ag-0 triggers EDS1-dependent immune signaling in response to the bacterial pathogen effector HopBA1 (Nishimura et al., 2017); overexpression of several other TIR-only genes also induces EDS1-dependent signaling (Bayless and Nishimura, 2020; Meyers et al., 2002; Nandety et al., 2013; Nishimura et al., 2017; Santamaria et al., 2019; Staal et al., 2008; Zhao et al., 2015), indicating that TIR-containing members share a conserved signaling pathway to mediate immune responses. Cell death mediated by TIR-containing proteins depends on the putative catalytic glutamate, which is conserved in ∼90% of plant TIR proteins (Wan et al., 2019). TIR NADase activity is essential but not sufficient for cell death and defense activation (Duxbury et al., 2020; Horsefield et al., 2019; Wan et al., 2019). Supporting this idea, estradiol-inducible expression of the *Pseudomonas syringae pv. tomato* effector AvrRps4 in transgenic Arabidopsis, which is recognized by the paired TNLs RRS1 and RPS4, fail to induce cell death but still up-regulate defense genes (Ngou et al., 2020). These results suggest that additional signaling components are needed to fully activate TIR-mediated immune responses.

In the current study, we found that plant TIR domain proteins had the activity of producing 2′,3′-cAMP/cGMP via hydrolysis of RNA/DNA. Nuclease activity was not sufficient for TIR-mediated cell death because TIR mutants retaining both nuclease and NADase activity but lacking the 2′,3′-cAMP/cGMP synthetase activity failed to trigger cell death in *Nicotiana benthamiana* (*N. benthamiana*). The structural basis of TIR proteins as bifunctional enzymes was revealed by cryo-EM structures of the L7 TIR domain (L7^TIR^) in complex with dsDNA. Structure-based mutagenesis analyses identified TIR mutations that specifically disrupted the 2′,3′-cAMP/cGMP synthetase activity *in vitro* greatly suppressed TIR-mediated cell death in *N. benthamiana*, indicating that this enzymatic activity is required for TIR signaling. Moreover, we found that *At*NUDT7 and the oomycete pathogen effector Avr3b function as 2′,3′-cAMP/cGMP phosphodiesterases (PDE) *in vitro*. Further evidence supporting a critical role for 2′,3′-cAMP/cGMP in TIR-mediated signaling, co-expression of wild-type (WT) *AtNUDT7*, but not its catalytic mutant, suppressed the cell death activity of RBA1 in *N. benthamiana*. As Nudix hydrolases are highly conserved throughout all organisms (McLennan, 2006), our findings open opportunities for the study of 2′,3′-cNMPs beyond plants.

## RESULTS

### TIR proteins exhibit nuclease and 2′,3′-cAMP/cGMP synthetase activity *in vitro*

When we examined the NADase activity of the TIR protein RBA1, we found that in the presence of Mg^2+^, RBA1 hydrolyzed many metabolites besides NAD^+^, such as ATP and GTP (Figures S1A and S1B). A shared feature of these *in vitro* RBA1 substrates is that they all contain phosphodiester bonds. This prompted us to test if DNA and RNA, whose backbones consist of phosphodiester bonds, are substrates of RBA1. As anticipated, the purified RBA1 protein exhibited activity of degrading PCR product (Figure 1A, left), plasmid DNA (Figure 1A, right) or *Arabidopsis* genomic DNA (gDNA) (Figure 1B). RBA1 displayed a much higher nuclease activity when *Arabidopsis* total RNA was used as substrate (Figures 1A-C, S1C and S1D). The nuclease activity appears to be a general feature of TIR proteins, because several TIR proteins tested including L7^TIR^ also displayed activity of degrading *Arabidopsis* gDNA and total RNA, although the activity varied among the proteins (Figures 1B and 1C). To further verify the nuclease activity of the RBA1 protein, we investigated whether it cleaves a DNA probe that generates a fluorescent product. Further supporting the data above, RBA1 and several other TIR proteins exhibited clear cleavage activity toward the DNA probe, albeit lower than that of the control DNase I (Figure 1D). Interestingly, unlike DNase I, TIR-catalyzed production of the fluorescent product appears not to obey Michaelis-Menten kinetics.

**Figure 1.**
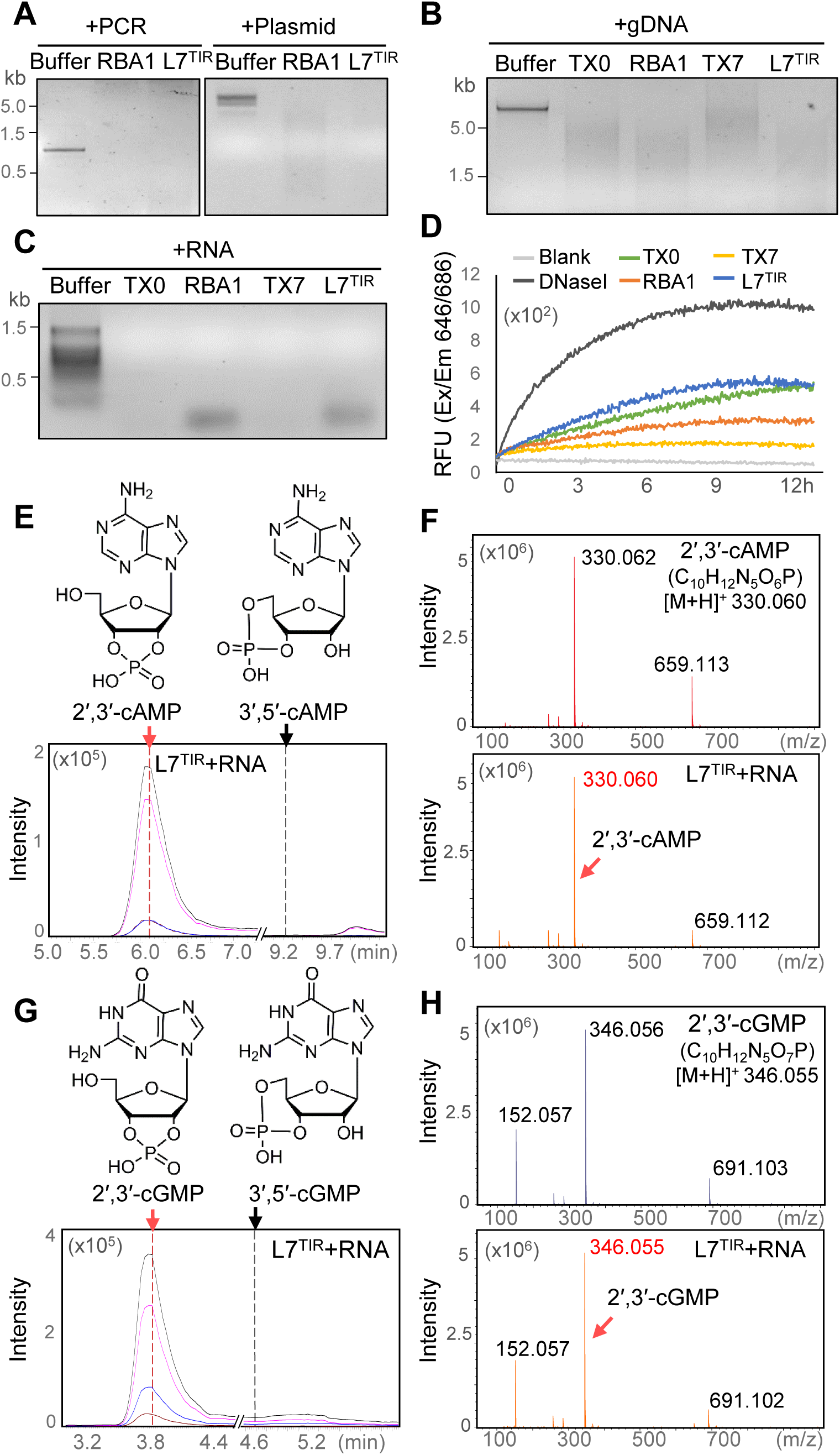
TIR proteins catalyze production of 2′,3′-cAMP/cGMP using RNA/DNA as substrates. (A-C) TIRs have DNase and RNase activity *in vitro*. TIR proteins indicated were incubated with 100ng PCR product (A, left pane), plasmid (A, right panel), *Arabidopsis* genomic DNA (gDNA) (B) and total RNA (C). After incubation of at 25°C for 16 hr, the reaction mixtures were analyzed by agarose gel electrophoresis. L:DNA ladder (Thermo Scientific 11581625). (D) TIR proteins display DNase activity toward a fluroscent probe. TIR proteins indicated were individually incubated with a fluroscent DNA probe. The fluroscent product yielded was meaured at 646/686nm (Extinction/Emission) by Microplate Reader SynergyH1 (BioTek) at different time points. (E-H) L7^TIR^ has 2′,3′-cAMP/cGMP synthetase activity using RNA as the substrate. Top of (E) and (G) : Molecular structures of 2′,3′-cAMP/3′,5′-cAMP and 2′,3′-cGMP/3′,5′-cGMP, respectively. Bottom of (E) and (G): L7^TIR^ protein was incubated with *Arabidopsis* total RNA at 25°C for 16 hr and the reaction products were analysed by LC-MS in MRM (multiple reaction monitoring) mode. A representative chromatogram of MRM analyses is shown for 2′,3′-cAMP (E) and 2′,3′-cGMP (G). MRM and retention time of 2′,3′-cAMP/cGMP and 3′,5′-cAMP/cGMP were confirmed using authentic standards. Red and black arrows indicate the positions of standard 2′,3′-cAMP/cGMP and 3′,5′-cAMP/cGMP, respectively. Black line indicates the total ion chromatogram (TIC), purple, blue and brown lines indicate three different transition 330.00>136.20, 330.00>99.15, 330.00>119.15 for 2′,3′-cAMP, and 346.00>152.15, 346.00>135.20, 346.00>110.15 for 2′,3′-cGMP, respectively. 2,3 (F) and (H): Comparison of the high-resolution MS spectra of the standard 2′,3′-cAMP (F) or 2′,3′-cGMP (H) and the L7^TIR^-generated products. 2′,3′-cAMP theoretical [M+H]^+^: 330.0597, standard [M+H]^+^: 330.0615, assay product [M+H]^+^: 330.0600. 2′,3′-cGMP theoretical [M+H]^+^: 346.0547, standard [M+H]^+^: 346.0558, assay product [M+H]^+^: 346.0552.

We then investigated whether the nuclease activity is associated with TIR-mediated cell death *in planta*. Cys132 of L7^TIR^ is highly conserved among plant TIR proteins and adjacent to the NADase catalytic residue Glu135 (Figure S1E). The equivalent cysteine residue has been shown to be required for cell death mediated by the TIR domain of grape TNL RPV1 (Williams et al., 2016) and L6^TIR^ (with 95% sequence identity with L7^TIR^) (Bernoux et al., 2011) in *N. tabacum* due to an unknown mechanism. Intriguingly, the residue at this position is substituted with Thr in the bacterial *A. baumannii* TIR (AbTIR) (Figure S1E), which possesses NADase activity but has no activity of inducing cell death in *N. tabacum* (Duxbury et al., 2020). These results suggest that the phenotype caused by mutation of the conserved cysteine residue in L6^TIR^ and RPV1^TIR^ *in planta* is probably not due to the absence of their NADase activity. Indeed, L7^TIR^ C132A had little impact on the NADase activity of L7^TIR^ (Figure S1F). Unexpectedly, however, the L7^TIR^ mutant protein also retained nuclease activity comparable to that of WT protein (Figure S1F). Together, these results suggest that the nuclease activity, if important, is not sufficient for the cell death activity of TIRs *in planta*.

If the nuclease activity *per se* is insufficient for TIR-mediated cell death, a plausible explanation for the data above is that there exist unknown molecules generated by TIR-mediated degradation of RNA/DNA for the cell death activity. To probe this possibility, we analyzed products of *Arabidopsis* total RNA or gDNA incubated with the L7^TIR^ protein using liquid chromatography-mass spectrometry (LC-MS). The results showed that incubation of L7^TIR^ with total RNA or gDNA but not L7^TIR^, the RNA, or DNA alone gave rise to two pronounced peaks, one with parent ion of 330 z/m (Figures 1E, 1F, S1G and S1H) and the other 346 z/m (Figures 1G, 1H, S1G and S1H), which correspond to the molecular weight of 3′,5′-cAMP and 3′,5′-cGMP, respectively. However, the retention time of these two unknown substances did not match to that of either standard 3′,5′-cAMP or standard 3′,5′-cGMP (Figures 1E, 1G and S1G). We speculated that the unknown substances are regioisomers of 3′,5′-cAMP and 3′,5′-cGMP, likely 2′,3′-cAMP and 2′,3′-cGMP. To test this possibility, we assayed standard samples of 2′,3′-cAMP and 2′,3′-cGMP by LC-MS. Indeed, the retention time of the first (330 z/m) and the second (346 z/m) peaks of the products generated by L7^TIR^ was identical to that of the standard 2′,3′-cAMP and 2′,3′-cGMP, respectively (Figures 1E, 1G and S1G). The identities of the two unknown substances were further confirmed by MRM (multiple reaction monitoring) (Figures 1E and 1G) and high-resolution MS analyses (Figures 1F and 1H). Similar results were obtained with the products from RBA1 incubated with *Arabidopsis* total RNA (Figure S1I). The 2′,3′-cAMP/cGMP synthetase activity of L7^TIR^ appears to be a conserved feature of TIR proteins, as several *Arabidopsis* TIR proteins, including RBA1, produced these two cNMPs in a similar manner when *Arabidopsis* gDNA (Figure S1K) or total RNA (Figure S1L) was used as substrate. As a control, fungal RNase T1, but not DNase I, showed similar activity in producing 2′,3′-cAMP/cGMP (Figure S1J). Taken together, our data demonstrate that TIR domains or TIR-only proteins act as 2′,3′-cAMP/cGMP synthetases with RNA/DNA as substrates.

### Cryo-EM structure of L7^TIR^ bound by dsDNA

While the NADase activity of TIR proteins is well documented (Horsefield et al., 2019; Ma et al., 2020; Wan et al., 2019), our data showed that TIR proteins have both NADase and 2′,3′-cNMP synthetase activity. To understand the mechanism of TIRs as bifunctional enzymes, we purified proteins using gel filtration to assay their enzymatic activity. We found that several TIR proteins, including L7^TIR^, were eluted at the void position of a Superose 6 gel filtration column, suggesting the formation of large ‘aggregates’ of these TIR proteins. The ‘aggregates’ appeared to contain nucleic acids as evidenced by UV absorbance at the wavelength of 260 nm (Figures S2A and S2B). Visualization of the negatively stained large molecular weight species revealed filaments or filament-like structures (Figure S2A). Notably, the filaments formed by L7^TIR^ following removal of the malt-binding protein (MBP) tag were more uniform (Figure S2A), rendering them amenable to structural analysis. We collected cryo-EM images of L7^TIR^ filaments (Figures S2C and S2D). After iterative 2D classification, helical segments converged into dozens of distinct classes with noticeable differences in the relative axial positioning of L7^TIR^ domains (Figures S2E and S2F). By grouping the classes with similar conformations, we were able to obtain distinct structures representing different states of DNA/RNA hydrolysis. We solved three distinct helical complex structures representing an initial DNA/RNA binding state, an intermediate state and an end state of DNA/RNA hydrolysis, with resolution of 3.3 Å, 2.7 Å and 2.6 Å, respectively (Figures S2G-J, Table S1).

In the structure of the initial dsDNA/RNA binding state, the L7^TIR^ filament is composed of 4 protofilaments with the two outer ones being made of L7^TIR^ (Figure 2A). The two L7^TIR^ protofilaments are not in direct contacts with each other. Sandwiched between the two L7^TIR^ protofilaments are two thinner protofilaments, which clearly come from dsDNA or dsRNA (Figure 2A), although the density does not allow a distinction. dsDNA was modeled to the helical-shaped density because it was more resistant to degradation by TIR proteins compared to RNA *in vitro* (Figures 1A, 1B, S1C and S1D). The lack of Mg^2+^ during protein purification might slow down the cleavage of the bound dsDNA by the L7^TIR^ protein. Nonetheless, dsDNA cleavage can be found in part of the L7^TIR^-dsDNA complex, as represented by the structures of the intermediate and the end state of L7^TIR^ filaments (Figures S2I-K). Interactions of the two L7^TIR^ protofilaments mediated by the two dsDNA protofilaments result in the formation of a superhelical structure (Figures 2A and S2N).

**Figure 2.**
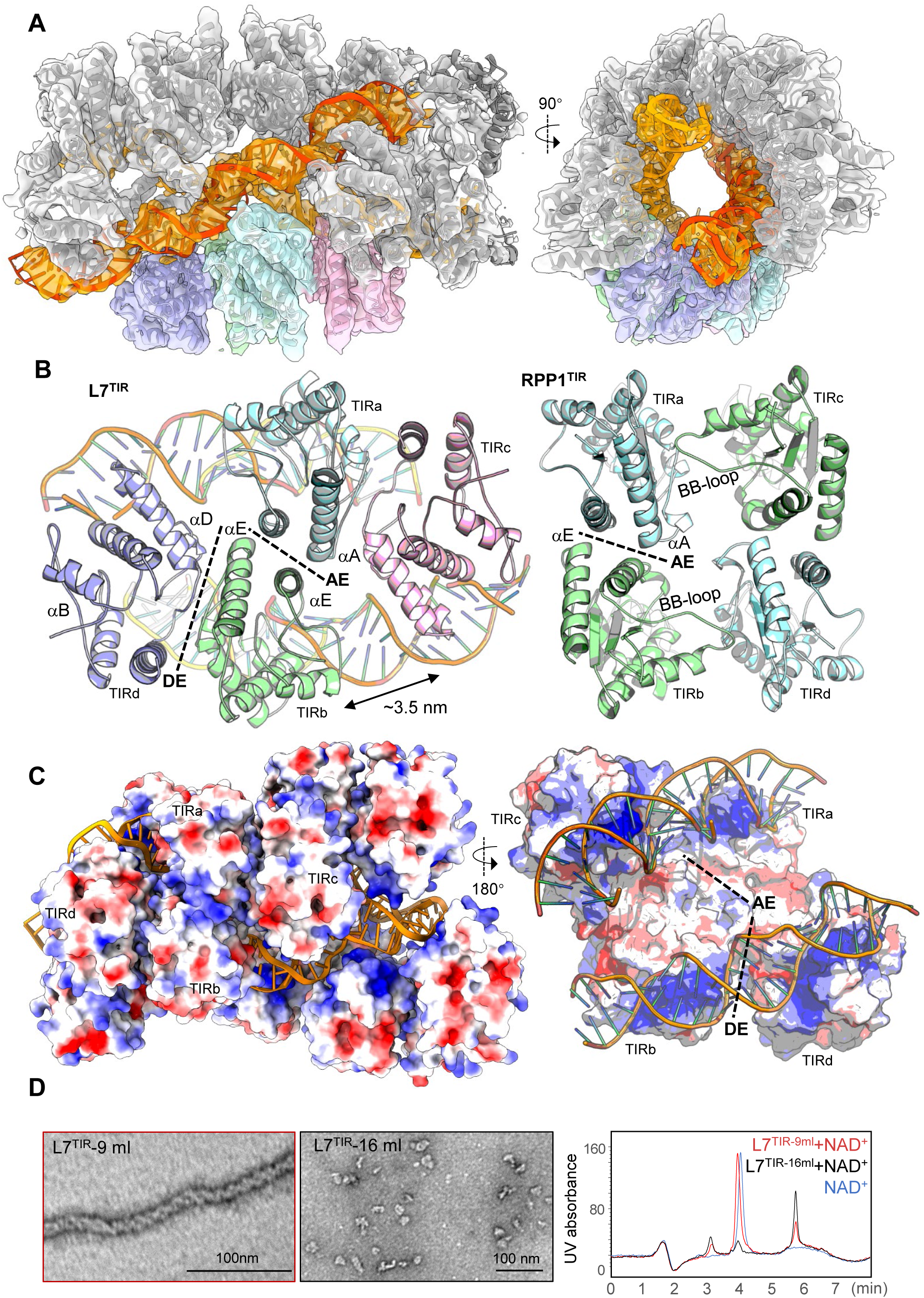
Cryo-EM structure of L7^TIR^ in complex with dsDNA. (A) Different views of the final cryo-EM map of L7^TIR^ filaments at an initial dsDNA-binding state. The cryo-EM density of L7^TIR^ and dsDNA is shown in gray and orange/orange red, respectively. Four L7^TIR^ domains described in (B) are highlighted by different colors. L7^TIR^ domains are shown in cartoon. (B) Left: Packing of L7^TIR^ domains (in different colors) in the cryo-EM structure of (A). Some structural elements are labeled. Dashed lines indicate L7^TIR^ dimerization interfaces. The middle two L7^TIR^ domains form symmetric dimers mediated by AE interface. Right: Packing of the four RPP1^TIR^ domains in the RPP1 resistosome (PDB code: 7CRC). The two BB-loops are indicated. (C) Different views of the cryo-EM structure of (A) shown in electrostatic surfaces. Left: Overall structure of L7^TIR^ filaments shown in electrostatic surface. Right: Longitudinal cut of the dsDNA-L7^TIR^ electrostatic interface. AE and DE interface are indicated by dashed lines. Red, blue and white represent negative, positive and neutral surfaces, respectively. The backbones of dsDNA are shown in orange. (D) dsDNA-bound L7^TIR^ has little NADase activity. Left and Middle: Negative staining images of L7^TIR^ (tag free) eluted at 9 ml and 16 ml in Superose 6 gel filtration column, respectively. Right: NADase activity of L7^TIR^ protein eluted at different elution volumes. 100μM NAD^+^ was incubated with 20 μM protein eluted at 9 ml (red) or 16 ml (black) in the presence of 10mM Mg^2+^ and products were analyzed with HPLC. Horizontal axis: Retention time of the samples (min). Vertical aixs: UV absorbance at 260 nM.

Two types of symmetric homodimers are discernible in the L7^TIR^ protofilaments. One is mediated by the AE interface (Figure 2B), which has been found in both crystal structures of TIR domains (Bernoux et al., 2011; Horsefield et al., 2019; Williams et al., 2014; Williams et al., 2016; Zhang et al., 2017) and cryo-EM structures of TNL resistosomes (Ma et al., 2020; Martin et al., 2020). The functional significance of this type of TIR dimer is well established (Ma et al., 2020; Martin et al., 2020; Nimma et al., 2017). The AE dimer as a protomer propagates to form the two L7^TIR^ protofilaments. The propagation of the AE dimers occurs through the interaction of two L7^TIR^ monomers of two adjacent AE dimers, resulting in the formation of the second type of L7^TIR^ dimer (Figure 2B). This L7^TIR^ dimer is nearly identical to the dimer in the crystal structure of L6^TIR^ mediated by the DE interface (Figure S2L), which is conserved in many TIR proteins and required for their cell death activity in *Nicotiana* species (Nishimura et al., 2017; Zhang et al., 2017). Interestingly, the L7^TIR^ protofilaments are reminiscent of a structural model of L6^TIR^ domain higher-order assembly (Zhang et al., 2017). Parallel packing of the two AE dimers via the DE interface generates a tetrameric L7^TIR^, which is strikingly different from the closed tetrameric TIR in the RPP1 resistosome (Figure 2B). Asymmetric L7^TIR^ dimers similar to those important for the NADase activity of the TNL resistosomes are absent in the tetrameric L7^TIR^. Along each L7^TIR^ protofilament are two positively charged surfaces between which the two dsDNAs are sandwiched (Figure 2C). We cannot assign the sequence of the modeled dsDNA due to its limited quality of cryo-EM density. However, the L7^TIR^ domains do not make base-specific contacts with the dsDNA, indicating non-sequence-specific binding to dsDNA/dsRNA by L7^TIR^. This is consistent with sequence-independent cleavage of *Arabidopsis* or barley total RNA *in vitro* (Figure S1C).

Formation of asymmetric TIR dimers is known to be required for NADase activity of TNLs (Ma et al., 2020; Martin et al., 2020). Such dimers are not present in the L7^TIR^ filament structure, suggesting that the L7^TIR^ oligomers cannot hydrolyze NAD^+^. To test this hypothesis, we assayed NADase activity of the L7^TIR^ protein eluted at the void position after gel filtration (Figure 2D, left) with high performance liquid chromatography (HPLC). As predicted, the L7^TIR^ ‘aggregate’ protein exhibited little activity in hydrolyzing NAD^+^ (Figure 2D, right). By contrast, the L7^TIR^ protein at the position with lower molecular weight species (Figure 2D, middle) nearly completely consumed NAD^+^ substrate under the same conditions. These results suggest that L7^TIR^ forms structurally distinct oligomers for its 2′,3′-cAMP/cGMP synthetase and NADase activity.

### Nucleic acid binding, AE and DE interfaces are important for 2′,3′-cAMP/cGMP synthetase activity of L7^TIR^

The distance between two adjacent L7^TIRs^ along the two outer protofilaments is ∼3.5 nm (Figure 2B), which is close to the length of one turn of dsDNA (3.4 nm). Such positioning allows the αD-helix of L7^TIR^ from one side of the outer protofilaments to periodically interact with the major groove of the dsDNA via electrostatic interaction (Figures 2A and 3A), which is characteristic for major groove binders of dsDNA (Eckel et al., 2003). A cluster of basic residues from this α-helix are solvent-exposed and largely conserved among TIR domain proteins (Figure S1E). Additionally, the loop connecting αB and β B (called BB-loop) of L7^TIR^ also contacts the backbones of dsDNA (Figure 3A). Notably, the conformation of the loop is different from that of its equivalent in the crystal structure of the L6^TIR^ domain (Figure S3A). BB-loops are also critical to mediate formation of asymmetric dimerization of RPP1^TIR^ (Ma et al., 2020) and Roq1^TIR^ (Martin et al., 2020). These results suggest that the BB-loop is likely important for both NADase and 2′,3′-cAMP/cGMP synthetase activity of TIR proteins. The AE interface in L7^TIR^ filaments is highly conserved in other TIRs (Figure S2L). As in the crystal structure of L6^TIR^ (Bernoux et al., 2011), the DE interface in the L7^TIR^ filaments is mainly mediated by van der Waals and hydrophobic contacts (Figure 3B).

**Figure 3.**
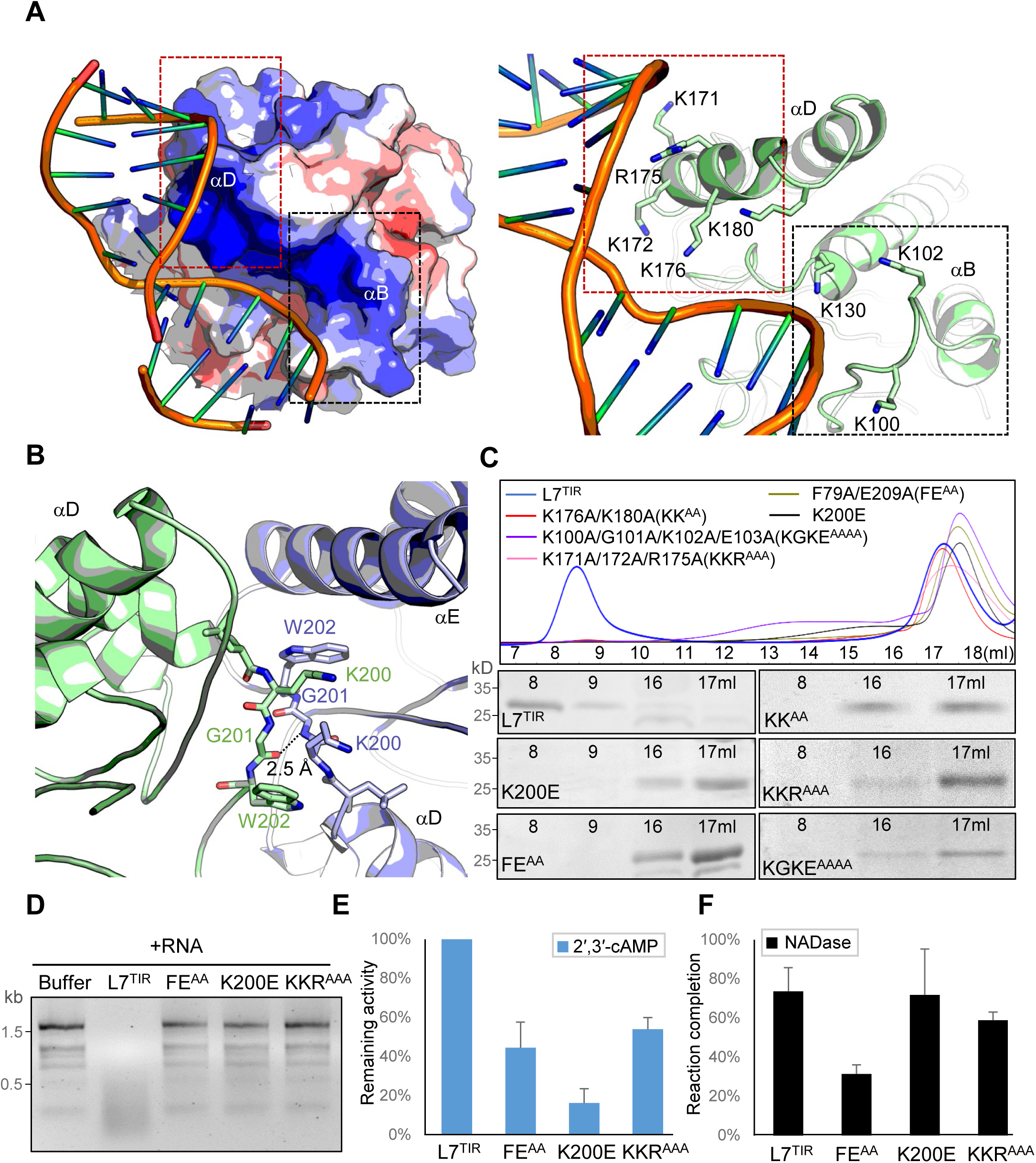
DNA binding and dimerization are required for 2′,3′-cAMP/cGMP synthetase activity of L7^TIR^. (A) Left: A close-up view of interaction between αD helix of L7^TIR^ and dsDNA, which are shown in eletrostatic surface and cartoon, respectively. Right: Detailed interactions of αD (red dashed frame) and the BB-loop (black dashed frame) with dsDNA. Some structural elements are labeled. (B) A close-up view of detailed interactions of the DE interface. (C) Gel filtration analyses of WT L7^TIR^ and L7^TIR^ mutants. Top and bottom: Gel filtration profiles and SDS-PAGE analyses of WT and L7^TIR^ mutant proteins, respectively. Horizontal axis: Elution volumes (ml) on Superose 6 column. FE^AA^, K200E and KKR^AAA^ are from the AE, DE and dsDNA binding interfaces of L7^TIR^ filaments, respectively. (D-F) Effect of L7^TIR^ mutations on nuclease (D), 2′,3′-cAMP synthetase (E), and NADase(F) activity of L7^TIR^. The assays for the nuclease and 2′,3′-cAMP synthetase activity were performed as described Figure 1C (6 hr reaction) and Figure 1E, respectively. Remaining activity was calculated as [MS intensity (area) of each sample/MS intensity (area) of WT L7^TIR^]×100%. The 2′,3′-cAMP synthetase activity of WT L7^TIR^ was normalized to 100% To assay NADase actvitiy, proteins indicated were individually incubated with NAD^+^ at 25°C for 16 hr and the unhydrolzyed NAD^+^ was quantified with HPLC. Reaction completion (%) of each sample was calculated as [1-(concentration of unhydrolyzed NAD^+^)/(concentration of NAD^+^ before reaction)]×100%.

To investigate whether the dsDNA-αD interaction is important for the 2′,3′-cAMP/cGMP synthetase activity of L7^TIR^, we made a simultaneous mutation K171A/K172A/R175A (KKR^AAA^) for the basic residues from this helix. In contrast to WT L7^TIR^, the KKR^AAA^ protein was eluted only at the position of the low molecular species during gel filtration (Figure 3C), indicating that the mutant has lost its filament-forming activity. Similar effects were also observed with K176A/K180A (KK^AA^) and K100A/G101A/K102A/E103A (KGKE^AAAA^) mutations predicted to disrupt αD and BB-loop interaction with dsDNA, respectively (Figure 3C). Furthermore, KKR^AAA^ significantly diminished both nuclease (Figures 3D and S3D) and 2′,3′-cAMP/cGMP synthetase (Figures 3E, S3B and S3C) activity of L7^TIR^. These basic residues are not involved in the formation of the AE dimer and the predicted asymmetric L7^TIR^ dimer, suggesting that they are dispensable to the NADase activity of the TIR protein. Indeed, the KKR^AAA^ mutation had little impact on the NADase activity of the TIR protein (Figure 3F).

The DE interface has been shown to be important for immune signaling of several TIRs (Nishimura et al., 2017; Zhang et al., 2017). We investigated whether the interface is required for the 2′,3′-cAMP/cGMP synthetase activity of L7^TIR^. Supporting an essential role of the DE interface in the filament-forming activity of L7^TIR^, an L7^TIR^ mutant with Lys200 at this interface mutated to Glu (Figure 3B) was eluted only at the low molecular species position (Figure 3C). As expected, the mutation significantly compromised the nuclease (Figures 3D and S3D) and synthetase (Figures 3E and S3C) activity of L7^TIR^ but had no detectable effect on the NADase (Figure 3F) activity of the TIR protein, thereby phenocopying the mutation of the dsDNA-binding residues. By comparison, all three L7^TIR^ enzymatic activities were greatly reduced by the mutation of residues (F79A/E209A, FE^AA^) from the AE interface (Figures 3C-F). Taken together, our structural and biochemical data showed that TIR oligomers mediated by different interface combinations confer 2′,3′-cAMP/cGMP synthetase or NADase activity.

### TIR proteins generate no 2′,3′-cyclophosphate-terminated RNA oligonucleotides

In addition to 2′,3′-cNMPs, RNA cleavage by RNases can also generate 2′,3′-cyclophosphate-terminated RNA oligonucleotides (Jackson, 2017). In fact, this type of RNA oligonucleotides generated by RNase T2 have been shown to act as PAMPs recognized by Toll-like receptor 8 (TLR8) to stimulate innate responses in human cells (Greulich et al., 2019; Ostendorf et al., 2020). Since TIR proteins have the activity to catalyze production of 2′,3′-cAMP/cGMP with RNA as substrate, we investigated the possibility of TIRs producing 2′,3′-cyclophosphate-terminated RNA oligonucleotides. Therefore, we assayed the products of *Arabidopsis* total RNA incubated with L7^TIR^ by LC-MS. Since 2′,3′-cAMP/cGMP are the major 2′,3′-cNMPs, we detected 2′,3′-cAMP- or 2′,3′-cGMP-terminated RNA dinucleotides. The results showed that the products contained no chemical species with molecular weight corresponding to any of these dinucleotides (Figure 4A). As a control, incubation of *Arabidopsis* total RNA with fungal RNase T1, a guanine base-specific RNase (Yoshida, 2001) strongly promoted the production of compounds with molecular weight corresponding to UGp and CGp (Gp: 2′,3′-cGMP). These results suggest that TIR proteins, as ribonucleases, catalyze the production of 2′,3′-cNMPs but not 2′,3′-cyclophosphate-terminated RNA oligonucleotides.

**Figure 4.**
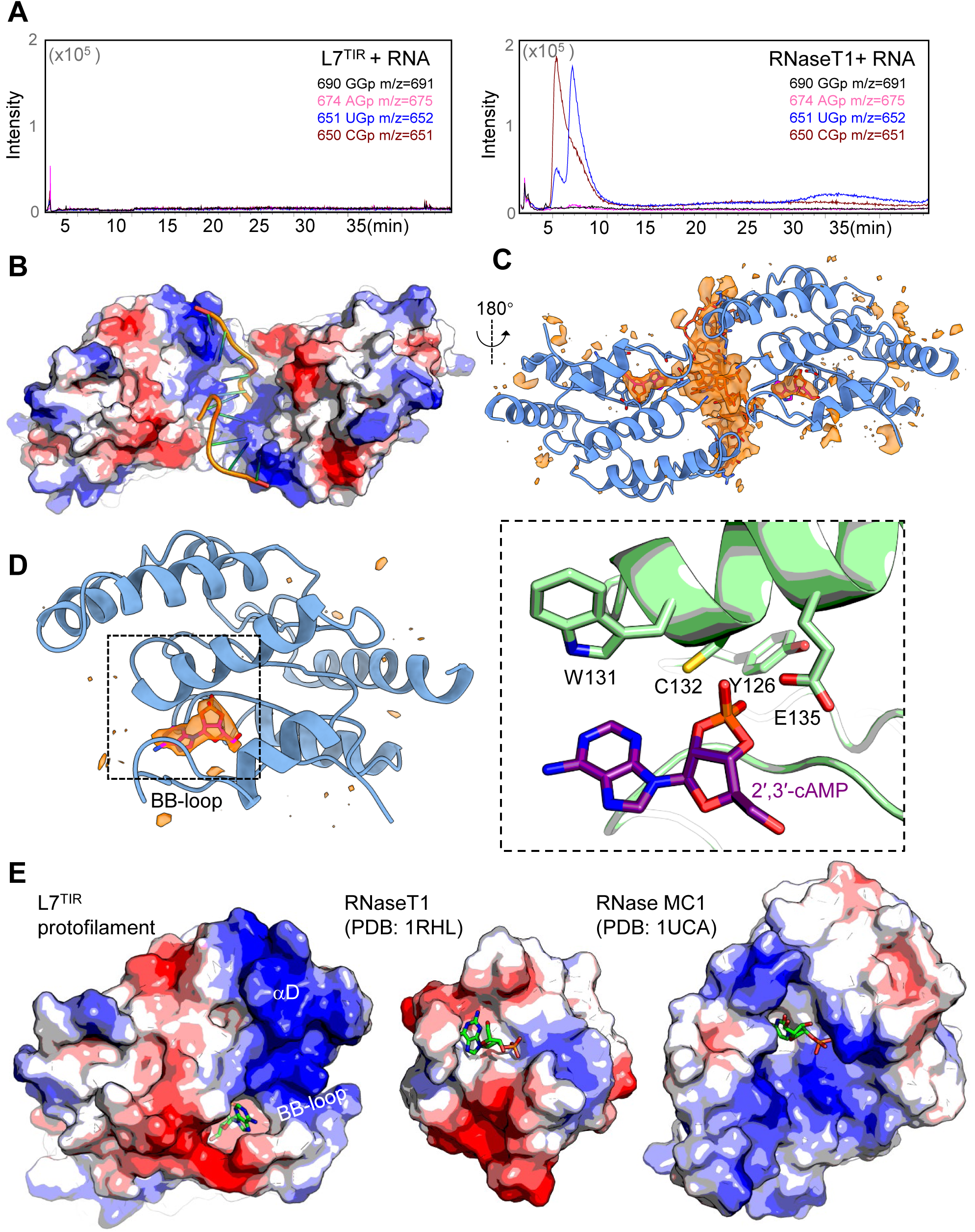
Hyrdolysis of RNA by L7^TIR^ produces no 2′,3′-cyclophosphate-terminated RNA oligonucleotides. (A) Analysis of RNA hydrolyzed products by L7^TIR^ (left) and fungal RNaseT1 (right) by LC-MS. *Arabidopsis* total RNA was incubated with RNaseT1 or L7^TIR^ at 25°C for 16 hr and reaction products were quantified by LC-MS. (B) Interaction of distorted ssDNA (cartoon) with L7^TIR^ (electrostatic surface). (C) Cryo-EM density (orange) not accounted by L7^TIR^ domain in the 3D reconstruction of L7^TIR^ filaments at the end state. Distorted ssDNA was built into the density. One 2′3′-cyclic AMP was built into the NAD^+^-binding pocket of each each L7^TIR^, in order to annotate two significant blobs of the different density. (D) Left: Cryo-EM density (orange) within the active site of L7^TIR^. Right: Detailed interactions of the modeled 2′,3′-cAMP with L7^TIR^. (E) Structural comparison of L7^TIR^ (left), RNaseT1 (middle, PDB code: 1RHL) RNase MC1(right, PDB code:1UCA). 2′,3′-cAMP bound by L7^TIR^ and 2′-GMP, 2′-UMP bound by RNaseT1. RNase MC1 are shown in stick.

The cryo-EM structure of L7^TIR^ filaments in the end state revealed that packing of the tetrameric TIR domains (Figure 2B, left) is similar to that in the initial and intermediate state (Figure S4A). Compared to the other two states, the end state displays less conformational heterogeneity, yielding the best resolution density (Figure S2H). A masked local refinement of two centrally located TIR domains allowed us to build a detailed atomic model of TIR domain with most of the side chains being easily identified (Figures 4B and S4B). No clear density for dsDNA is present in the end state and nucleic acid hydrolysis appears to be completed (Figure 4C). There is an extra density blob inside the predicted NAD^+^-binding pocket of L7^TIR^ roughly with the size of a single nucleotide (Figure 4C). Interestingly, the conformation of the BB-loop of L7^TIR^ is nearly identical with that in the crystal structure of RUN1^TIR^ bound by NADP^+^ and bis-Tris [Bis(2-hydroxyethyl) amino-tris (hydroxymethyl) methane] (Figure S4C). The quality of the density doesn’t allow us to unambiguously determine the identity of the molecule bound in the pocket, but docking of nucleotides and nucleotide derivatives showed that a 2′,3′-cNMP (2′,3′-cAMP modeled) best fits the topology of the density (Figure 4D). The modeled 2′,3′-cAMP completely overlaps with bis-Tris in the structure of RUN1^TIR^ (Figure S4B) and forms interactions with the side chains of Trp131, Cys132 and Glu135 (Figure 4D). The modeled 2′,3′-cyclic NMP is buried in the pocket (Figure 4D, left). By comparison, the active sites of RNase T1 (Ishikawa et al., 1996) and RNase MC1 (Numata et al., 2003) are largely solvent-exposed, making it possible for them to bind an oligonucleotide for cyclization of the 3′-phosphate group (Figure 4E). The smaller size of the binding pocket of L7^TIR^ can only allow binding of a mononucleotide for cyclization reaction. These structural features can explain why L7^TIR^ does not produce di- or probably any other longer 2′,3′-cyclophosphate-terminated RNA oligonucleotides.

### 2′,3′-cAMP/cGMP synthetase activity is required for TIR-mediated cell death

Our data showed that TIR proteins are bifunctional enzymes with both NADase and 2′,3′-cAMP/cGMP synthetase activity. This raised the question whether the 2′,3′-cAMP/cGMP synthetase activity is required for TIR signaling. While the conserved Cys132 was dispensable for the NADase and nuclease activity of L7^TIR^ (Figure S1F), the equivalent residue is important for the cell death activity of L6^TIR^ (Bernoux et al., 2011) and RPV1^TIR^ (Williams et al., 2016) in *N. tabacum*. These results suggest that this conserved residue may be specifically important for the 2′,3′-cAMP/cGMP synthetase activity of TIR proteins. Indeed, C132A abrogated much of the 2′,3′-cAMP/cGMP synthetase activity of L7^TIR^ (Figure S5A). To validate this result with another TIR protein, we mutated the equivalent residue Cys83 of RBA1 to Ala and then examined nuclease, 2′,3′-cAMP/cGMP synthetase and NADase activity of the mutant protein. As anticipated, C83A greatly impaired the 2′,3′-cAMP/cGMP synthetase (Figures 5A and S5B) but had no detectable effect on the NADase (Figure 5B) and a modest effect on the nuclease (Figure S5C) activity of RBA1. Like mutations of the equivalent residues of L6^TIR^ and RPV1^TIR^, C83A nearly completely suppressed the cell death phenotype of RBA1 in *N. benthamiana* (Figure 5C). Together, these results support an essential and conserved role of 2′,3′-cAMP/cGMP synthetase activity in TIR-mediated cell death, confirming the idea that nuclease activity is not sufficient for TIR-mediated cell death.

**Figure 5.**
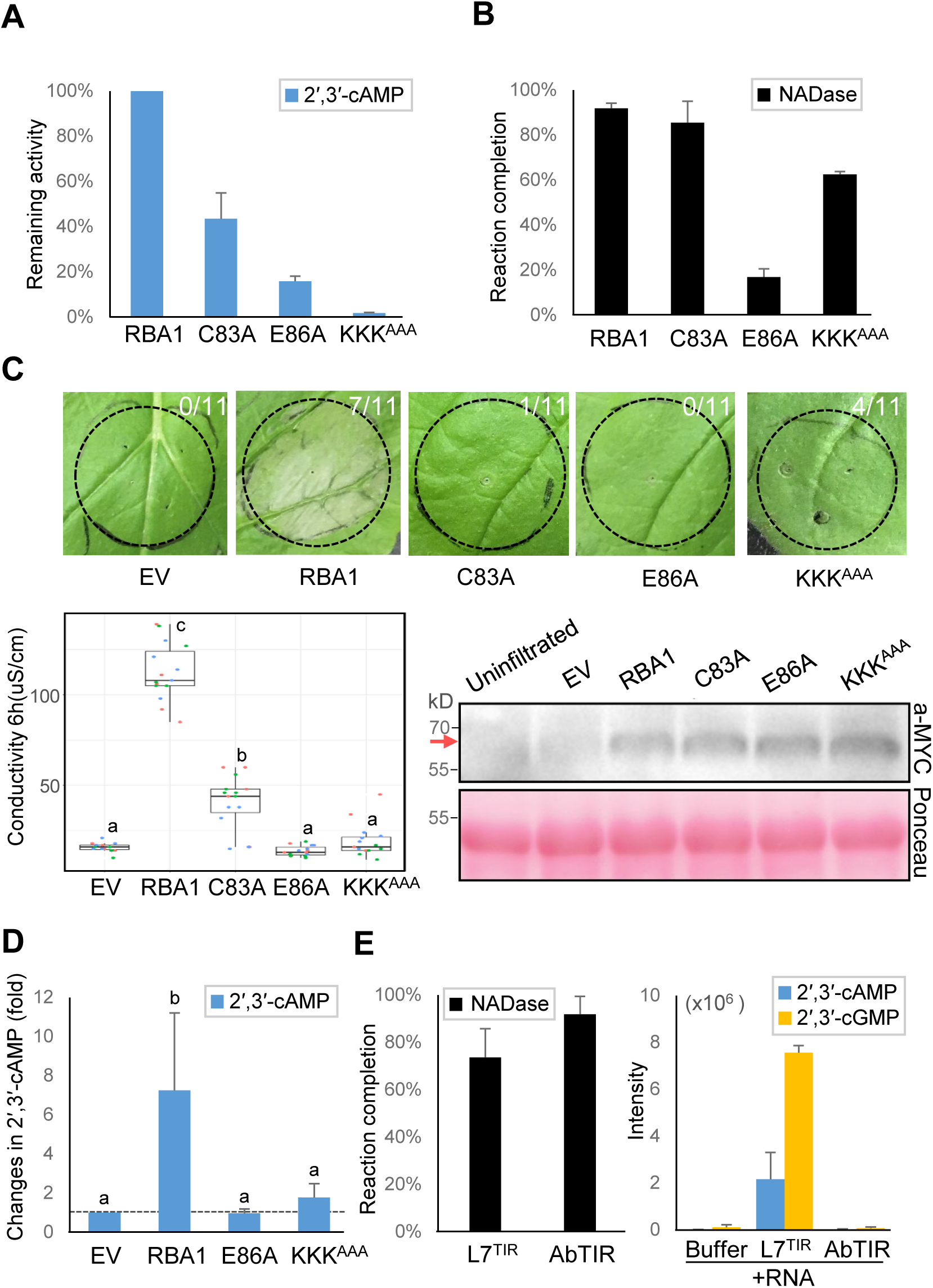
2′,3′-cAMP/cGMP synthetase activity is required for RBA1-mediated cell death. (A and B) 2′,3′-cAMP synthetase (left) and NADase (right) activity of WT and RBA1 mutant proteins. Vertical axis: MS intensity (area) of 2′,3′-cAMP produced by mutant protein compared with WT RBA1 protein (A) and percentage of NAD^+^ consumed (B). The 2′,3′-cAMP synthetase activity of WT RBA1 was normalized to 100%. The assays for 2′,3′-cAMP synthetase and NADase activity of different RBA1 proteins were performed as described in Figure 3E and Figure 3F, respectively. (C) Cell death phenotype of *N. benthamiana* plants transiently expressing WT and *RBA1* mutants. Top: Cell death was visually assessed and photographed at 4 dpi. The numbers indicate the numbers of leaves displaying cell death out of the total number of leaves infiltrated. Bottom left: Cell death was quantified by the electrolyte leakage assay 3 days post infection (dpi). The plants were infiltrated in the same conditions as those for the electrolyte leakage assays. Colors indicate bilogical replicates. Significance was calculated with Tukey′s HSD test (n = 15, α = 0.05; shared lowercase letters indicate no significant difference). Bottm right: protein blots for WT and RBA1 mutants. Total protein was extracted from 2 dpi leaves and subjected to immunoblot using antibodies indicated. Ponceau staining is shown to indicate loading. Red arrow points the expected size of RBA1. (D) 2′,3′-cAMP levels in *N. benthamiana* plants expressing WT and *RBA1* mutants. Leaf extract of *N. benthamiana* plants expressing indicated constructs was analyzed by LC-MS. Vertical axis: MS intensity of 2′,3′-cAMP of each construct. For quantification, internal standard 8-Br-2′,3′-cAMP was used. Significance was calculated with Tukey′s HSD test (n = 3, α = 0.05; shared lowercase letters indicate no significant difference). The level of 2′,3′-cAMP in *N. benthamiana* expressing EV was normalized to 1.0. (E) AbTIR has NADase but no detectable 2′,3′-cAMP/cGMP synthetase activity. NADase (left), 2′,3′-cAMP/cGMP synthetase activity (right) of L7^TIR^ and AbTIR were performed as described in Figure 3F and Figure 3E, respectively.

Like the mutation of basic residues in L7^TIR^ (Figures 3F-H), a simultaneous mutation K122A/K123A/K130A (KKK^AAA^) of the equivalent residues from helix αD in RBA1 nearly abrogated the 2′,3′-cAMP/cGMP synthetase (Figures 5A and S5C) and nuclease (Figure S5B) but only slightly impacted the NADase (Figure 5B) activity of RBA1. As a control, the catalytic mutation E86A resulted in loss of the three enzymatic activities of RBA1 (Figures 5A, 5B S5B and S5C). We then took advantage of RBA1 KKK^AAA^ to further confirm whether the 2′,3′-cAMP/cGMP synthetase activity is important for TIR-mediated cell death. The cell death phenotype caused by RBA1 in *N. benthamiana* was completely suppressed by the catalytic E86A mutation that eliminated the NADase activity of RBA1 (Figures 5B and 5C), further confirming an essential role for the NADase enzymatic activity in TIR signaling. Importantly, the cell death phenotype was also substantially suppressed by the KKK^AAA^ mutation (Figure 5C), indicating that the 2′,3′-cAMP/cGMP synthetase activity is required for TIR-mediated signaling. Together, our data support a critical and conserved role for the cluster of basic residues from αD in the cell death activity of TIRs.

To further verify the significance of 2′,3′-cAMP/cGMP in TIR signaling, we investigated if expression of a TIR protein promotes accumulation of 2′,3′-cAMP/cGMP *in planta*. We assayed levels of these two non-canonical 2′,3′-cNMPs in *RBA1*-expressing *N. benthamiana* plants by LC-MS. Compared to empty vector (EV), expression of WT *RBA1* significantly enhanced accumulation of both 2′,3′-cAMP and 2′,3′-cGMP (Figures 5D and S5D). In contrast, *N. benthamiana* plants expressing *RBA1* catalytic mutant E86A had much lower levels of 2′,3′-cAMP/cGMP comparable to those of EV-expressing plants. Similarly, KKK^AAA^ with reduced RBA1-mediated cell death also greatly reduced the accumulation of 2′,3′-cAMP/cGMP. Together, these results indicate that RBA1-mediated cell death in *N. benthamiana* is accompanied by increased 2′,3′-cAMP/cGMP levels.

AbTIR has NADase activity (Essuman et al., 2018) but fails to induce cell death when expressed in *N. tabacum* (Duxbury et al., 2020). Sequence alignment indicated that the cluster of basic residues conserved in plant TIR proteins are not conserved in AbTIR (Figure S1E). Given an important role of these basic residues in catalyzing production of 2′,3′-cAMP/cGMP by RBA1 (Figures 5A and S5C) and L7^TIR^ (Figures 3E and S3C), this result suggests that AbTIR lacks 2′,3′-cAMP/cGMP synthetase activity. Indeed, while AbTIR had comparable NADase activity to L7^TIR^ as previously reported (Duxbury et al., 2020), the bacterial TIR protein displayed no nuclease and 2′,3′-cAMP/cGMP synthetase activity (Figures 5E and S5E). These results may explain why AbTIR has no cell death activity *in planta* and provide additional evidence for a critical role of 2′,3′-cAMP/cGMP synthetase activity in TIR-mediated cell death.

### 2′,3′-cAMP/cGMP phosphodiesterases inhibit TIR-mediated cell death

Our results support a crucial role of 2′,3′-cAMP/cGMP in TIR-mediated cell death. Conceivably, intracellular homeostasis of these noncanonical cNMPs has to be maintained in the absence of biotic or abiotic stress conditions. Although *in vivo* substrates of *At*NUDT6,7 remain unknown, the catalytic residue Glu154 is required for *At*NUDT7 inhibition of EDS1 signaling (Ge et al., 2007). These results prompted us to hypothesize that *At*NUDT6,7 act as 2′,3′-cAMP/cGMP PDEs to negatively regulate EDS1-dependent signaling. To test this hypothesis, we evaluated 2′,3′-cAMP/cGMP-hydrolyzing activity of recombinant *At*NUDT7 protein purified from *E. coli* by HPLC. In support of our hypothesis, *At*NUDT7 exhibited PDE activity toward 2′,3′-cAMP/cGMP but not their positional isomers 3′,5′-cAMP/cGMP *in vitro* (Figures 6A, S6A and S6B). Furthermore, substitution of the catalytic residue Glu154 with Gln resulted in loss of the 2′,3′-cAMP/cGMP PDE activity (Figures 6A and S6A). These results demonstrate that *At*NUDT7 acts as a 2′,3′-cAMP/cGMP PDE *in vitro*. Further supporting this conclusion, the *At*NUDT7 protein hydrolyzed 2′,3′-cAMP/cGMP produced by L7^TIR^ but had no effect on L7^TIR^-mediated degradation of *Arabidopsis* total RNA (Figures S6A and S6C).

**Figure 6.**
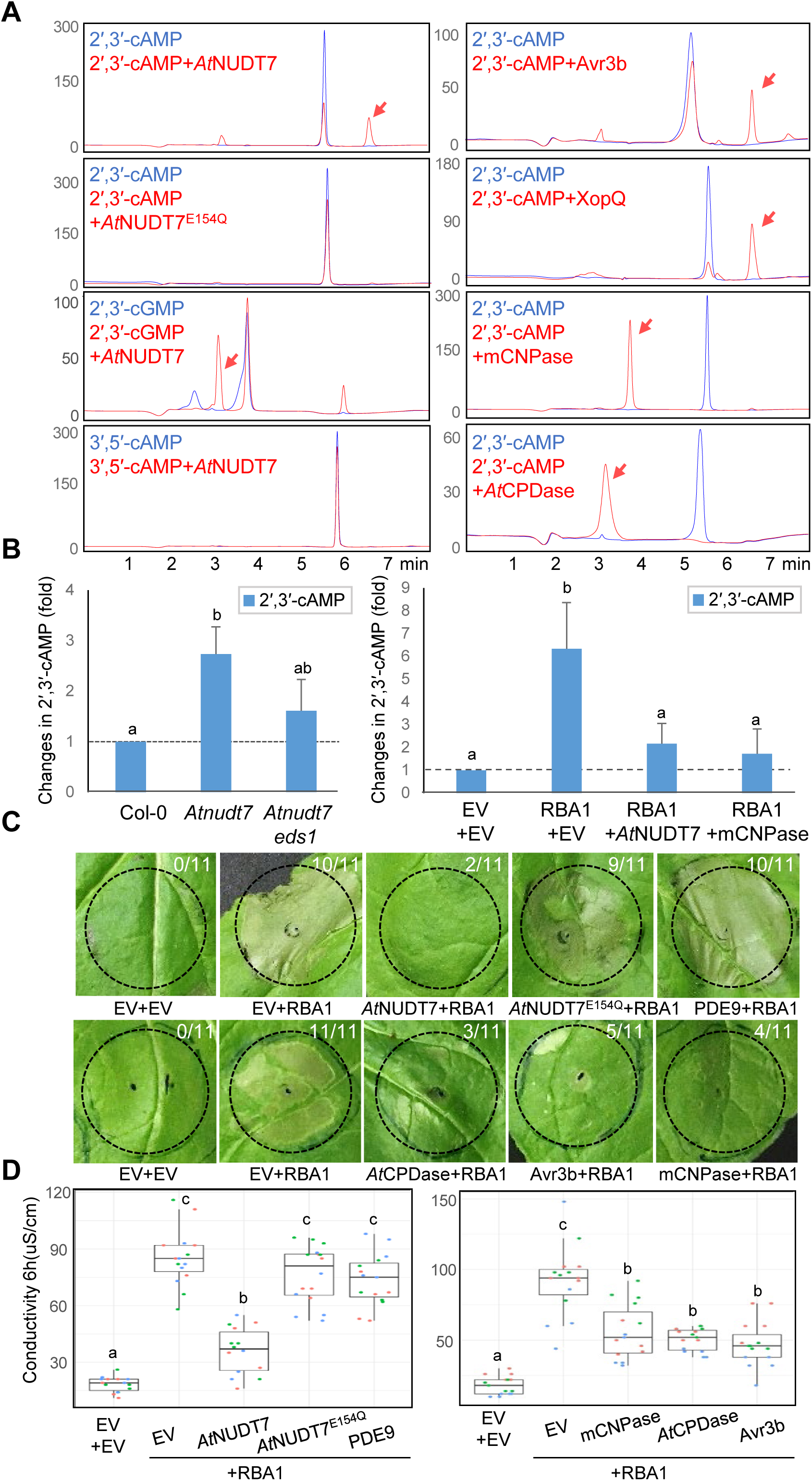
2′,3′-cAMP/cGMP phosphodiesterases suppress RBA1-mediated cell death. (A) Hydrolysis of 2′,3′-cAMP/cGMP by proteins indicated. Purified enzymes indicated were incubated with 2′,3′-cAMP (100 μM) or 2′,3′-cGMP (100 μM) in the presence of Mg^2+^ (10.0 mM) at 25°C for 16 h. Mixture of each reactions was then centrifuged and the supernatant was analyzed with HPLC. Peaks from absorbance of the hydrolyzed products of 2′,3′-cAMP/cGMP are indicated by red arrows. Vertical axis: UV absorbance at 260 nM. (B) 2′,3′-cAMP/cGMP phosphodiesterases reduce the levels of 2′,3′-cAMP *in planta*. Left: *Atnudt7* displys a higher level of 2′,3′-cAMP than WT plants. Leaf extract of 4 weeks old *Arabidopsis* plants indicated was analyzed by LC-MS as described in Figure 5D. The level of 2′,3′-cAMP in *Arabidopsis* Col-0 was normalized to 1.0. Right: Expression of 2′,3′-cAMP/cGMP phosphodiesterases reduces the 2′,3′-cAMP levels of *N. benthamiana* plants expressing *RBA1*. Empty vector (EV) or constructs indicated were co-expressed with *RBA1* in *N. benthamiana* plants and leaf extract was analyzed by LC-MS as described in Figure 5D. The level of 2′,3′-cAMP in *N. benthamiana* expressing EV+EV was normalized to 1.0. For quantification, internal standard 8-Br-2′3′cAMP was used. Significance was calculated with Tukey′s HSD test (n = 3, α = 0.05; shared lowercase letters indicate no significant difference). (C) Co-expression with *AtNUDT7* (top panel) or other 2′,3′-cAMP/cGMP phosphodiesterases (bottom panel) supress RBA1-mediated cell death in *N. benthamiana*. Empty vector (EV) or constructs indicated were co-expressed with *RBA1* in *N. benthamiana* plants. Cell death was visually assessed and photographed at 4 dpi. The assays were performed as described in Figure 5C. (D) Ion leakage assay of *N. benthamiana* plants transiently co-expressing *RBA1* with different constructs indicated. The assays were performed as described in Figure 5C.

We next investigated whether the 2′,3′-cAMP/cGMP PDE activity is required for *AtNUDT7* inhibition of EDS1 signaling. To this end, we first quantified the 2′,3′-cAMP/cGMP levels of *Atnudt7* plants. In further support of previous data (Bartsch et al., 2006; Ge et al., 2007), *Atnudt7 Arabidopsis* plants exhibited a stunted growth phenotype (Figure S6D). Importantly, levels of both 2′,3′-cAMP (Figure 6B, left) and 2′,3′-cGMP (Figure S6E, left) were markedly elevated in the mutant plants as compared to those of WT plants, suggesting that the absence of *At*NUDT7 reduces hydrolysis of these cNMPs. Interestingly, while not statistically significant, the levels of 2’, 3’-cAMP/cGMP in *Atnudt7 eds1* double mutant plants appear to be higher than those of WT plants (Figures 6B and S6E). This result suggests an EDS1-dependent and an EDS1-independent role in promoting the production of these cNMPs. These results support *At*NUDT7 as a 2′,3′-cAMP/cGMP PDE *in vivo* and reinforce our conclusion that 2′,3′-cAMP/cGMP synthetase activity is required for TIR-mediated and EDS1-dependent signaling.

If the 2′,3′-cAMP/cGMP PDE activity of *At*NUDT7 is important for inhibition of EDS1 signaling, then *AtNUDT7* is predicted to block TIR-mediated cell death. To test this hypothesis, we assessed the effect of *AtNUDT7* on RBA1-mediated cell death in *N. benthamiana*. As anticipated, co-expression with *AtNUDT7* substantially suppressed the cell death activity of RBA1 (Figures 6C, 6D and S6F). The cell death suppressing activity of *AtNUDT7* was nearly abrogated by the catalytic mutation E154Q. These results demonstrate that the inhibition of RBA1-mediated cell death by *AtNUDT7* is dependent on its 2′,3′-cAMP/cGMP PDE activity. In further support of this conclusion, 2′,3′-cAMP/cGMP levels were significantly reduced in *N. benthamiana* leaves co-expressing *RBA1* and *AtNUDT7* than in leaves co-expressing *RBA1* and EV (Figures 6B, right and S6E, right). Unlike *AtNUDT7*, co-expression with 3′,5′-cGMP-specific *PDE9* (Hanna et al., 2012) had little effect on RBA1-mediated cell death in *N. benthamiana* (Figures 6C, 6D and S6F). Taken together, our data indicate that *AtNUDT7* suppresses RBA1-mediated cell death by acting as a 2′,3′-cAMP/cGMP PDE to downregulate the levels of 2′,3′-cAMP/cGMP, supporting a critical role for both of these cNMPs in TIR signaling.

We then asked whether 2′,3′-cNMP PDEs can generally inhibit RBA1-mediated cell death. 2′,3′-cyclic-nucleotide 3′-phosphodiesterase (CNPase) is an animal enzyme that hydrolyzes 2′,3′-cNMPs to their respective 2′-nucleotides (Myllykoski et al., 2016), although its biological function remains undefined. The enzymatic activity of mouse CNPase (mCNPase) was confirmed by our HPLC assay (Figures 6A and S6A). Like *At*NUDT7, mCNPase had no activity to hydrolyze 3′,5′-cAMP/cGMP *in vitro* (Figures S6A and S6B). Co-expression with *mCNPase* in *N. benthamiana* significantly compromised RBA1-mediated cell death (Figures 6C, 6D and S6F). Similar observations were made with the *Arabidopsis* 2′,3′-cyclic phosphodiesterase (*At*CPDase) (Figures 6A, 6C, 6D, S6A and S6F).

The function of 2′,3′-cAMP/cGMP in TIR-triggered and EDS1-dependent cell death suggests their critical role in immune signal transduction. If so, it is conceivable that phytopathogens have evolved strategies to disrupt this signaling mechanism. Our data presented above show that Nudix hydrolases can act as 2′,3′-cAMP/cGMP PDEs to suppress EDS1 signaling. Nudix hydrolases are also present in many pathogens such as bacteria, oomycetes and fungi, and some Nudix effectors have been found to be critical for pathogen virulence activity (Dong and Wang, 2016). Examples are the effector proteins *Xanthomonas euvesicatoria* XopQ (Adlung and Bonas, 2017) and *Phytophthora sojae* Avr3b (Kong et al., 2015). These results collectively suggest that these two effector proteins may have 2′,3′-cAMP/cGMP PDE activity. We purified the two effector proteins from *E. coli* and assayed their 2′,3′-cNMP PDE activity by HPLC. As hypothesized, both effector proteins displayed activity to hydrolyze 2′,3′-but not 3′,5′-cAMP/cGMP in the assay (Figures 6A, S6A and S6B). Furthermore, co-expression with *Avr3b* notably compromised RBA1-mediated cell death in *N. benthamiana* (Figures 6C, 6D and S6F). A similar experiment was not performed for XopQ because the effector is recognized by the endogenous *N. benthamiana* TNL Roq1 (Schultink et al., 2017). Nonetheless, these results serve as further evidence for pathogen-mediated negative regulation of 2′,3′-cAMP/cGMP-induced signaling by 2′,3′-cAMP/cGMP PDEs during pathogen infection. Metabolism of 2′,3′-cAMP/cGMP by XopQ also provides an explanation for XopQ-mediated inhibition of EDS1-dependent cell death in *Nicotiana* species (Adlung and Bonas, 2017).

### Model on the role of 2′,3′-cAMP/cGMP in TIR signaling

Having shown that downregulation of 2′,3′-cAMP/cGMP levels suppresses RBA1-mediated cell death, we next asked whether upregulation of these two cNMPs promotes the cell death activity of RBA1. Treatment of *N. benthamiana* plants with 8-Br-2′,3′-cAMP, a cell-permeable analogue of 2′,3′-cAMP, had no detectable effect on the cell death activity of RBA1, likely because the analogue is unable to mimic all functions of 2′,3′-cAMP in plants or/and both 2′,3′-cAMP and 2′,3′-cGMP are needed to promote RBA1 signaling. We then tested if the HopBA1 effector-induced cell death in *Arabidopsis* Ag-0 is accompanied with increased levels of 2′,3′-cAMP/cGMP. Consistent with a previous study, delivery of *HopBA1* by *Pseudomonas fluorescens* Pf0-1 in leaves induced a macroscopically visible cell death in *Arabidopsis* Ag-0 at 48 hpi (Figure S7A). LC-MS analysis showed that 2′,3′-cAMP/cGMP levels in these plants were higher than those in EV carrying Pf0-1 infiltrated plants (Figure 7A). Together with previous data (Nishimura et al., 2017), these results indicate a positive correlation between 2′,3′-cAMP/cGMP levels and RBA1-mediated immune signaling.

**Figure 7.**
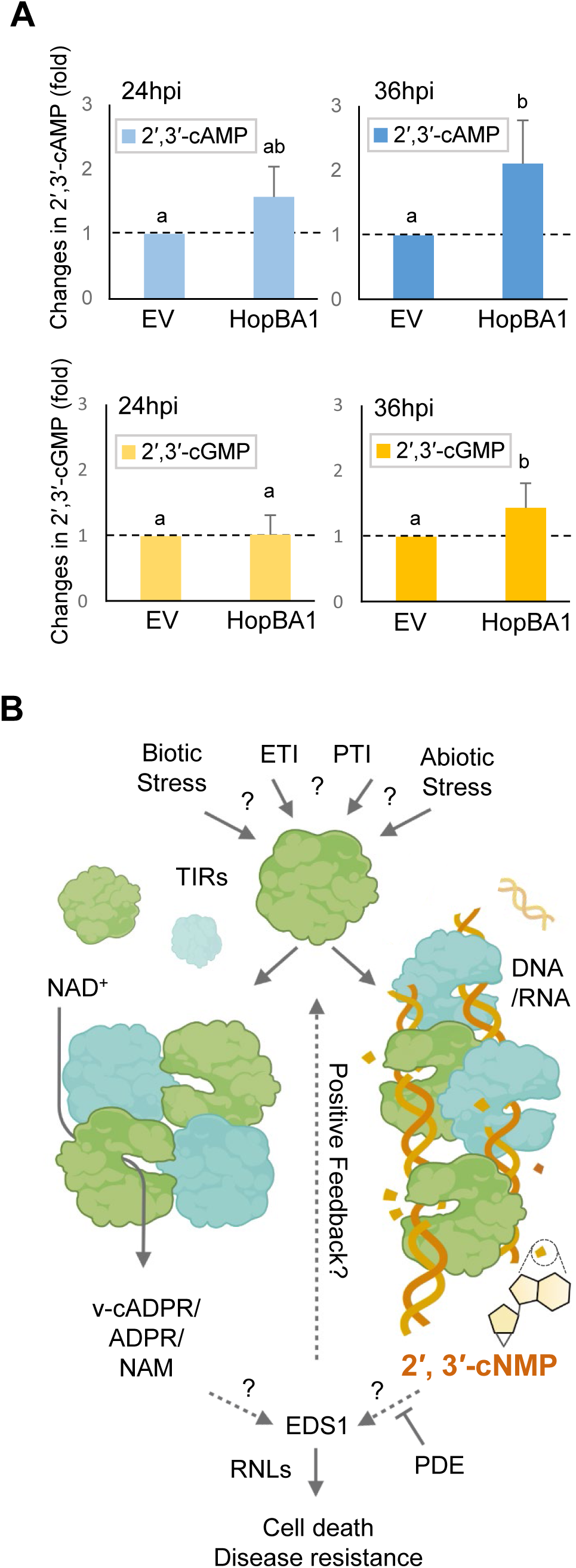
Working model on TIR-mediated signaling. (A) 2′,3′-cAMP (top) and 2′,3′-cGMP (bottom) levels are enhanced by the bacterial effector HopBA1 in *Arabidopsis* Ag-0 plants. Pf0-1 carrying HopBA1 was infiltrated into *Arabidopsis* accession Ag-0. Leaf extract was analyzed by LC-MS at 24 hpi (left) and 36 (right) hpi. The levels of 2′,3′-cAMP/cGMP extracted from EV carrying Pf0-1 infiltrated leaves at each time point were normalized to 1.0. (B) Model on signaling mechanism of 2′,3′-cAMP/cGMP. Monomeric TIR domain is shown in light green and cyan. In response to different stimuli, TIR domain proteins can form oligomers for NADase and 2′,3′-cAMP/cGMP synthetase activity. Products such as v-cADPR from the NADase activity and 2′,3′-cAMP/cGMP act together to trigger initiation of EDS1-dependent immune signaling.

Our results suggest that production of 2′,3′-cNMPs by TIRs is upstream of EDS1. Unexpectedly, however, expression of *RBA1* in *eds1* mutants of *N. benthamiana* induced no pronounced accumulation of 2′,3′-cAMP/cGMP (Figure S7B). This is consistent with the observation that 2′,3′-cAMP/cGMP levels in *Atnudt7 eds1* plants were comparable with those of WT *Arabidopsis* (Figures 6B and S6E). However, downregulation of these two cNMPs by 2′,3′-cAMP/cGMP PDEs significantly compromised EDS1-dependent immune signaling in *N. benthamiana* (Figure 6). A model that could reconcile these data is the existence of positive feedback between TIR-catalyzed 2′,3′-cAMP/cGMP and activation of EDS1-dependent signaling, which is negatively regulated by 2′,3′-cAMP/cGMP PDEs (Figure 7B). The model agrees with the observation that the strength of TIR-mediated HR-like cell death is strongly correlated with protein levels (Zhang et al., 2004). Since TIR signaling is activated during PTI (Pruitt et al., 2021; Roth et al., 2017; Tian et al., 2020), this model may also explain why PTI activation potentiates HR cell death in transgenic *Arabidopsis* plants expressing AvrRps4 (Ngou et al., 2020) and flg22 enhances HR-like cell death mediated by AtTN3 in *N. benthamiana* (Nandety et al., 2013). Thus, 2′,3′-cAMP/cGMP-promoted EDS1 signaling appears to involve a self-amplification mechanism.

## DISCUSSION

2′,3′-cNMPs are intermediates of mRNA turnover by RNases. Enzymes engaging in specific production of these noncanonical cNMPs remained unidentified. Due to the very limited number of studies directed at them, little is known about the biology of these noncanonical cNMPs. In the current study, we provide biochemical and structural evidence that plant TIR proteins function not only as NADases but also as 2′,3′-cAMP/cGMP synthetases. TIR mutants lacking either enzymatic activity lost their cell death activity, indicating that the dual enzymatic activity is important for TIR function. Furthermore, down regulation of 2′,3′-cAMP/cGMP levels by 2′,3′-cyclic PDEs from different origins, including pathogens, suppressed TIR-mediated cell death; conversely, upregulation of 2′,3′-cAMP/cGMP in *Atnudt7* plants or the delivery of HopBA1 into RBA1-containing host cells was associated with cell death. These results collectively support an important role for the 2′,3′-cAMP/cGMP synthetase activity in TIR-mediated immune signaling. The present study not only identified a novel enzyme family for the production of 2′,3′-cNMPs but also established an essential role for them in mediating EDS1-dependent signaling, opening up new horizons for investigation of these noncanonical cNMPs.

### Catalytic mechanism of TIR

Although both DNA and RNA were hydrolyzed by TIR proteins to produce 2′,3′-cAMP/cGMP *in vitro*, whether DNA or RNA or both are substrates of TIR 2′,3′-cAMP/cGMP synthetases remains unknown. *In vitro*, RNA appears to be a much more favorable substrate. It is rather surprising that DNA acts as a substrate for the production of 2′,3′-cAMP/cGMP since it does not have a 2′-hydroxyl group. The mechanism of TIR-catalyzed production of 2′,3′-cNMPs remains to be defined. Nonetheless, the observation that mutations of the conserved cysteine residue abrogated 2′,3′-cAMP/cGMP synthetase but not nuclease activity suggests a two-step mechanism for production of these cNMPs, in which cleavage of RNA/DNA may be the first step. A model of this may be that RNA/DNA cleavage results in flipping and binding of a nucleotide to the catalytic site of TIRs for the cyclization reaction. Consistently, a patch of density in the catalytic pocket of the intermediate state appears to come from a flipped nucleotide (Figure S2K). These data would explain why ATP and GTP were not substrates of TIRs for production of 2′,3′-cAMP/cGMP (Figure S7C). Thus, plant TIRs are different from the human DNase/RNase endonuclease G, which is a non-specific nuclease directly cleaving RNA/DNA for apoptotic cell death (Cregan et al., 2004). TIR proteins as nucleases appear to have different kinetics from DNase I (Figure 1D) probably due to the formation of filament- or filament-like structures.

### Mechanism of TIR proteins as bifunctional enzymes

In conjunction with previous studies, our biochemical data show that plant TIR proteins act as bifunctional enzymes with 2′,3′-cAMP/cGMP synthetase and NADase activity. In contrast with canonical bifunctional enzymes that generally harbor two structural domains with separate catalytic sites (Moore, 2004), the dual enzymatic activity of TIR proteins are encoded by the same structural domain. Structural and biochemical data showed that TIRs form different oligomers for their NADase and 2′,3′-cAMP/cGMP synthetase activities (Figures 2 and 3), thereby providing the structural basis of TIR proteins as bifunctional enzymes. As far as we are aware of, this represents an unprecedented mechanism of how dual enzymatic activity is encoded by a single structural domain. In addition to the AE and DE interfaces, RNA/DNA substrates clearly have an important role in forming TIR oligomers for 2′,3′-cAMP/cGMP synthetase activity, because mutations predicted to disrupt DNA binding activity resulted in loss of the enzymatic activity (Figure 3). While a TIR domain alone has NADase activity, assembly of TNL resistosomes mediated by the NBD domain of TNLs significantly promotes TIR NADase activity (Ma et al., 2020; Martin et al., 2020), indicating that NBD-assisted formation of the tetrameric TIR in the resistosome is important for this enzymatic activity.

AE and DE interfaces have been shown to be generally required for TIR-mediated immune signaling by current and previous data (Nishimura et al., 2017; Williams et al., 2016; Zhang et al., 2017). Our cryo-EM structure showed that formation of the L7^TIR^-dsDNA complex is mediated by both interfaces, revealing the mechanism underlying simultaneous requirement of the two interfaces for TIR functions. While the AE interface is highly conserved among TIRs, variations exist in the DE interface. These variations, as seen in the crystal structure of RPP1^TIR^, allow no propagation of the DE and AE interfaces to form a filament-like structure (Zhang et al., 2017). However, the RPP1^TIR^ protein was active in catalyzing production of 2′,3′-cAMP/cGMP albeit with lower activity (Figure S7D), suggesting that filament formation is not absolutely required for the 2′,3′-cAMP/cGMP synthetase activity of TIRs. This can be explained by our structural observation that a L7^TIR^ tetramer mediated by the AE and DE interfaces is sufficient for binding one complete turn of dsDNA/dsRNA (Figure 3A). This tetramer is incompatible with the tetrameric TIR that is important for the NADase activity (Ma et al., 2020; Martin et al., 2020), suggesting that 2′3,-cAMP/cGMP synthetase and NADase activity are exclusive in a given oligomeric TIR. Consistent with this conclusion, our biochemical data showed that the L7^TIR^-dsDNA complex had little NADase activity (Figure 2D). The nucleic acid-bound oligomeric TIR can function as a distinct form of resistosome, but further studies are needed to support this conclusion.

### 2′,3′-cNMPs-promoted self-amplification of EDS1 signaling

Both 2′,3′-cNMP synthetase and NADase activities are essential for RBA1-mediated HR cell death in *N. benthamiana*, but whether and how the two activities interact remains unknown. Triggering initiation of TIR immune signaling may also require the dual enzymatic activity. Once initiated, TIR signaling can promote production of 2′,3′-cNMPs, which in turn further amplify TIR signaling. Initiation of EDS1 signaling may cause alterations, for example in structures and/or stability of dsDNA/dsRNA, which allow a TIR protein to access the otherwise poor substrates. This model agrees with the observations that some TNLs require nuclear localization (Wirthmueller et al., 2007; Zhu et al., 2010) for function and NLR activation induces EDS1-dependent DNA damage (Rodriguez et al., 2018). Thus, TIRs can guard the integrity of RNA/DNA, perturbations of which by pathogen- or host-derived components may activate TIR-mediated immune signaling. This would provide an explanation of how TIR-only proteins sense the presence of pathogen effectors despite the lack of the C-terminal pathogen-sensing LRR domain as demonstrated in both CNLs (Wang et al., 2019b) and TNLs (Ma et al., 2020; Martin et al., 2020). Addressing of how dual enzymatic activity of the TIR domain is regulated between different TIR proteins would assist in understanding why only a few of ∼30 TIR-only genes in the *Arabidopsis* genome cause an HR cell death phenotype when overexpressed in *N. benthamiana* (Nandety et al., 2013). Clearly, more studies, including the identification of predicted 2′,3′-cAMP/cGMP receptors, are needed to unveil the signaling mechanism of these noncanonical cNMPs. These studies may also aid to determine whether these cNMPs act as second messengers or not.

### Negative regulation of 2′,3′-cAMP/cGMP levels

*AtNUDT7* is a negative regulator of EDS1 signaling (Bartsch et al., 2006; Ge et al., 2007), but the underlying biochemical mechanism remains elusive. Although several small molecules, such as ADPR, NADH and NAD were shown to be substrates of *At*NUDT7 Nudix hydrolase *in vitro*, changes in their levels were not detected in *Atnudt7* mutant plants (Ge et al., 2007). We found that 2′,3′-cAMP/cGMP but not their regioisomers 3′,5′-cAMP/cGMP were substrates of AtNUDT7 (Figures 6A and S6B). Consistently, accumulation of 2′,3′-cAMP (Figure 6B, left) and 2′,3′-cGMP (Figure S6E, left) were significantly enhanced in *Atnudt7* leaves, supporting a role of these two cNMPs in immune signaling. However, the phenotypes of *Atnudt7-1* plants are largely abolished under conditions of reduced stress (Straus et al., 2010), suggesting that 2′,3′-cNMPs alone are insufficient for EDS1 activation. This is consistent with the idea that both 2′,3′-cNMP synthetase and NADase activities are required for TIR-mediated signaling. Notably, the Nudix effector XopQ and Avr3b also exhibited 2′,3′-but not 3′,5′-cAMP/cGMP PDE activity *in vitro* with the latter effector suppressing RBA1-mediated cell death in *N. benthamiana* (Figure 6C). These results suggest that pathogens have evolved strategies for targeting 2′,3′-cAMP/cGMP-induced signaling to defeat plant immunity.

Nudix hydrolases belong to a highly conserved family of enzymes across all organisms and have broad substrate specificities (McLennan, 2006). Our finding that *At*NUDT7, XopQ and Avr3b act as 2′,3′-cAMP/cGMP PDEs opens opportunities for investigations of this enzyme family and the two noncanonical cNMPs beyond plants. In addition to Nudix hydrolases, plant 2′,3′-AMP/cGMP PDEs were also found in *Triticum aestivum* germ and shown to possess the ability to hydrolyze 2′,3′-cNMPs into 2′-NMPs (Tyc et al., 1987). Two homologues (At4g18930 and At4g18940) of this group of PDEs are present in *Arabidopsis*. Biochemical data showed that At4g18930 has similar enzymatic activity (Genschik et al., 1997), which is further confirmed by our enzymatic assays. It will be of interest to know whether these non-Nudix 2′,3′-cAMP/cGMP PDEs have role in regulating EDS1-dependent signaling. We wish to mention that only changes in 2′,3′-cNMPs levels in leaves were detected in our study. Given the diversities of TIR and Nudix hydrolase-encoding genes families in plant genomes, accumulation of these non-canonical cNMPs might be differentially regulated in other plant organs and tissues.

### Multiple potential functions of 2′,3′-cAMP/cGMP

Accumulation of RBA1 is not only induced by HopBA1, but is also broadly correlated with HR cell death triggered by NLRs (Nishimura et al., 2017), and a role of RBA1 in immune signaling mediated by the CNL ZAR1 has been shown recently (Martel et al., 2020). Consistent with a contribution of TIRs in CNL-mediated ETI, more recent data revealed a direct functional link between *At*TN13 and RPS5 in Arabidopsis (Cai et al., 2021). These results agree with gene expression profiling analysis revealing biotic stress-induced expression of TIR-only and TN genes (Nandety et al., 2013). In addition to ETI, accumulating evidence supports a role of TIR signaling in PTI responses (Pruitt et al., 2021; Roth et al., 2017; Tian et al., 2020). Consistent with this, EDS1-dependent signaling is required for basal resistance to virulent pathogens (Bonardi et al., 2011; Rietz et al., 2011). These results suggest a broad role for TIR signaling in ETI and PTI responses, which are intricately connected (Ngou et al., 2021; Pruitt et al., 2021; Tian et al., 2020; Yuan et al., 2021). A role of 2′,3′-cAMP/cGMP in plant abiotic stresses was suggested by recent data. Wounding (Van Damme et al., 2014), heat and dark stress conditions (Kosmacz et al., 2018) induce accumulation of cellular 2′,3′-cAMP/cGMP in plants. Moreover, 2′,3′-cAMP mediates stress granule formation (Kosmacz et al., 2018), which are ubiquitous, non–membrane-bound assemblies of protein and RNA inside cells, and mimics the abiotic response (Chodasiewicz et al., 2021) in *Arabidopsis*, strongly supporting its role in abiotic stress responses. Interestingly, *AtNUDT7* is one of the most induced genes in response to 2′,3′-cAMP (Chodasiewicz et al., 2021). *AtNUDT7* transcripts are responsive to numerous abiotic and biotic stress conditions (Straus et al., 2010), suggesting a broad role for *AtNUDT7* and possibly 2′,3′-cAMP/cGMP in the regulation of plant stress responses. However, it remains unknown whether TIR signaling is directly involved in abiotic stress responses induced by 2′,3′-cAMP in plants. Future investigations are needed to determine whether and how 2′,3′-cNMPs are involved in the interplay between PTI and ETI, TNL- and CNL-triggered, and abiotic-biotic responses. Answering these questions may provide new signaling paradigms for stress biology.

## ACKNOWLEDGMENTS

We thank Jian Shi, Yue Sun, Xiaoxiao Zhang for their advice and help with cryo-EM data collection. We thank Ruben Garrido Oter, Frederickson Entila and Yuang Wu for their advice with statistical analysis. We thank Sabine Metzger for her help on 2′,3′-cAMP and 2′,3′-cGMP confirmation of standard and assays. We thank Ulla Neumann for her advice and help on negative staining EM. We acknowledge the Tsinghua University Branch of the China National Center for Protein Sciences (Beijing) and National University of Singapore for providing the cryo-EM facility support and the computational facility support on the cluster of Bio-Computing Platform. We acknowledge the biocenter, University of Cologne, for providing MS facilities We acknowledge greenhouse faculty of Max Planck Institute for Plant Breeding Research for providing *N. benthamiana* plants. This work was supported by the Alexander von Humboldt Foundation (a Humboldt professorship to J.C.), the Max-Planck-Gesellschaft (a Max Planck fellowship to J.C.), Deutsche Forschungsgemeinschaft SFB-1403-414786233 (J.C. and P.S.-L.) and Germany′s Excellence Strategy CEPLAS (EXC-2048/1, Project 390686111) (J.C. and P.S.-L.) and Ministry of Health, Singapore, NMRC-OFIRG grant (MOH-000382-00 to W.B.).

## Figure legends

**Figure S1. TIR proteins have 2**′,**3**′**-cAMP/cGMP synthetase activity, related to Figure 1**

(A) SDS-PAGE analysis of TIR proteins indicated. MBP-GST tagged TX0, MBP-tagged TX7 and L7^TIR^ were purified from *E. coli*. SUMO-tagged RBA1 was purified from SF21 insect cells and the tag was removed by PreScission protease. Red arrows point the expected protein size. L: protein ladder (Thermo Scientific, 26617).

(B) Different *in vitro* substrates of RBA1. 100 μM ATP, GTP, CTP or TTP was incubated with 10 μM RBA1 supplemented with 10 mM Mg^2+^. After incubation at 25°C for 16 hr, reaction mixtures were centrifuged and analyzed by LC-MS. Vertical axis:percentage of samples hydrolyzed by RBA1.

(C) L7^TIR^ protein as a nuclease hydrolyzes total RNA from *Arabidopsis* and barley. 5 μM L7^TIR^ protein was incubated with *Arabidopsis* total RNA (AtRNA) or barley total RNA (HvRNA) for 3 hr and the samples were then visualized by agarose gel.

(D) L7^TIR^ protein displays lower nuclease activity when dsDNA is used as a substrate. The assay was perfromed as (C) except that a PCR product was used as a substrate. The products were visualized by agarose gel at different time points. Note that the PCR prodcut was not completely cleaved after 16 hr as compared to complete cleavage of plant total RNA after 3 hr shown in (C).

(E) Sequence alignment of TIR proteins. Residues at the AE interface are indicated by inverse blue triangles and basic residues in αD are indicated by black frames. The yellow asterisk on the top indicates the conserved cysteine residue.

(F) C132A mutation has little effect on L7^TIR^ NADase (left) and nuclease (middle) activity. Protein input is shown in the right panel.

(G) MRM analyses of 2′,3′-cAMP/cGMP (top) or 3′,5′-cAMP/cGMP (bottom) standards.

(H) L7^TIR^ protein, *Arabidopsis* total RNA and gDNA alone display negligible activity of producing 2′,3′-cAMP/cGMP. For comaprison, vertical axis is shown in the same level as that in Figure 1E and 1G.

(I) High resolution MS analyes of RBA1-RNA reaction products. The assay was performed as Figure 1E, F, G and H.

(J) RNaseT1 but not DNase I display 2′,3′-cAMP/cGMP synthetase activity. The assays were performed as described in Figure 1E and 1G.

(K and L) 2′,3′-cAMP/cGMP synthetase activity is conserved among TIRs. 2′,3′-cAMP/cGMP synthetase activity of TIR proteins indicated was assayed with *Arabidopsis* genomic DNA and total RNA as substrates. The assays were performed as described in Figure 1E and 1G. Vertical axis:MS intensity (area) of 2′,3′-cAMP/cGMP.

**Figure S2 cryo-EM structure of L7**^**TIR**^ **filaments, related to Figure 2**

(A) TIR proteins have activity of forming filaments *in vitro*. MBP-tagged L7^TIR^, L7^TIR^, L6^TIR^, RBA1 and TX0 after affinity purification were subjected to a Superose 6 gel filtration column. UV absorbance at 260 nM (A260) and 280 nM (A280) were shown by red and blue lines respectively. Fractions eluted at void position (9 mL, highlighted in red) revealed filaments or filament-like structures by negative staining EM as shown on the right side.

(B) UV measurement of purified WT L7^TIR^ and L7^TIR^ DNA binding mutant proteins (Figure3C). Horizontal axis: UV absorbance at different wavelengths. Note that only WT L7^TIR^ but not the DNA binding mutant showed a strong UV absorbance at A260, reflecting the nucleotide binding ability of L7^TIR^.

(C) A representative cryo-EM micrograph of L7^TIR^ filaments.

(D) Flowcharts for cryo-EM structure determination of L7^TIR^ filaments.

(E) Representative 2-D averages of L7^TIR^ filaments showing the existence of heterogeneous complexes. Red arrows indicate upper boundary of individual L7^TIR^ domain density. Different vertical positioning of L7^TIR^ suggests presence of multiple helical symmetries.

(F) Initial determination of helical parameters of L7^TIR^ filaments. Preliminary assignment of helical parameters was conducted using RELION 3.08, combining direct image analysis (left panel) and helical profile analysis (right panel). As shown in the helical profile plot, helical rise was determined to be ∼33.5Å. (512/20) *1.306 Å. Double peak is common in preliminary analysis of thin flexible filaments.

(G) Procedure for 3D reconstruction of L7^TIR^ filaments in an initial state

(H) Procedure for 3D reconstruction of L7^TIR^ filaments in an intermediate and an end state

(I) The final cryo-EM density maps for L7^TIR^ filaments in the initial (left), intermediate (middle) and end (right) state. The regions highlighted with the blue dashed lines indicate two L7^TIR^ domains. Pocket distance: Vertical distance between the densities of residue Ser129 of opposite L7^TIR^ measured in Chimera. This pair of L7^TIR^ bound to the same stretch of nucleic acids.

(J) The postprocessed cryo-EM density maps for L7^TIR^ filaments in the initial (left), intermediate (middle) and end (right) state. Cryo-EM density for dsDNA is shown in orange.

(K) Density not accounted by TIR domain is shown in orange in the intermediate state complex. Distorted double stranded DNA was built into the density to simulate the transition state.

(L) Structural alignment of TIR dimers mediated the DE interface (A) and AE interface (right).

(M) 3-repeat segment (left) and central repeat segment (right) FSC plot s at 0.143, 0.333, 0.5 in the final model of L7^TIR^ filament in the initial (top), intermediate (middle) and end (bottom) states.

(N) Left and middle: Different views of the cryo-EM structure of L7^TIR^ filaments shown in cartoon. Right: Schematic diagram of L7^TIR^ filaments cut along the orange piece of DNA in the middle panel. The sinusoids represent dsDNA.

**Figure S3. DNA-binding and dimerization are important for L7**^**TIR**^ **synthetase activity, related to Figure 3**

(A) Structural alignment between L7^TIR^ protomer and L6^TIR^ (3OZI). The BB-loop is indicated by the red arrow.

(B) Protein input of Figure 3D-F. SDS-PAGE analysis of MBP-tagged WT and L7^TIR^ mutant proteins. FE^AA^: F79A/E209A; KKR^AAA^: K171A/K172A/R175A.

(C) Effect of L7^TIR^ mutations on 2′,3′-cGMP synthetase activity. The assay was performed as described Figure3E.

(D) Effect of L7^TIR^ mutations on nuclease activity using gDNA as substrate. The assay was performed ad described in Figure 3D.

(E) A representative cryo-EM density of L7^TIR^ AE interface.

(F) A representative cryo-EM density of L7^TIR^ DE interface.

**Figure S4. Hyrdolysis of RNA by L7**^**TIR**^ **produces no 2**′,**3**′**-cyclophosphate-terminated RNA oligonucleotides, related to Figure 4**

(A) Organization of the L7^TIR^ domain tetramer shown in Figure 2B (left) is largely conserved in all three states. Shown is the initial state complex TIR tetramer (left) that fits into the intermediate state (middle) and end (right) state density maps.

(B) Structural alignment of L7^TIR^ with the crystal structure of RUN1^TIR^ in complex with NADP^+^ and bis-Tris.

(C) A representative cryo-EM density of the 2′,3′-cAMP binding pocket.

**Figure S5. 2**′,**3**′**-cAMP/cGMP synthetase activity is critical for RBA1 cell death activity, related to Figure 5**

(A) Cys132 is required for the 2′,3′-cAMP/cGMP synthetase activity of L7^TIR^. The 2′,3′-cAMP/cGMP synthetase activity of WT L7^TIR^ was normalized to 100%. The assays were performed as Figure 3E.

(B) 2′,3′-cGMP synthetase activity of WT and RBA1 mutant proteins. The assay was performed as Figure 5A.

(C) Left and middle:Nucelase activity of WT and RBA1 mutant proteins. The assays were performed as described in Figure 1B. Right: SDS-PAGE analysis of RBA1 and RBA1 mutant proteins. S80A: mutant at the potential dsDNA interacting residue. K149E: mutant at DE interface.

(D) 2′,3′-cGMP levels in *N. benthamiana* plants expressing WT and *RBA1* mutants. The level of 2′,3′-cGMP in *N. benthamiana* expressing EV was normalized to 1.0. For quantification, internal standard 8-Br-2′3′cAMP was used. Significance was calculated with Tukey′s HSD test (n = 3, α = 0.05; shared lowercase letters indicate no significant difference).

(E) Left: AbTIR displays negligible nuclease activity. L7^TIR^ was applied as positive control. Right: Protein input of AbTIR and L7^TIR^.

**Figure S6 2**′,**3**′**-cAMP/cGMP PDEs suppress TIR-mediated cell death, related to Figure 6**

(A) SDS-PAGE of proteins used in Figure 6A.

(B) *At*NUDT7, mCNPase, *At*CPDase, Avr3b and XopQ have no 3′,5′-cAMP/cGMP phosphodiesterase activity. Proteins shown in (A) were used to assay the phosphodiesterase activity of the proteins. The assays were conducted as described in Figure 6A.

(C) *At*NUDT7 and mCNPase hydrolyze 2′,3′-cAMP/cGMP produced by L7^TIR^ (bottom) but do not affect nuclease activity of L7^TIR^ (top). Assays for 2′,3′-cAMP/cGMP synthetase and nuclease activity of L7^TIR^ were conducted as described in Figure 1B and Figure 1J, respectively.

(D) Growth phenotypes of 4 weeks old *Arabidopsis* plants indicated, scale bar: 1 cm.

(E) Effect of 2′,3′-cAMP/cGMP phosphodiesterases on the level of 2′,3′-cGMP *in planta*. Left: *Atnudt7* displays a higher level of 2′,3′-cGMP than WT plants (left). Extract of *Arabidopsis* plants indicated was analyzed by LC-MS. The level of 2′,3′-cGMP in *Arabidopsis* Col-0 was normalized to 1.0. Right: Expression of 2′,3′-cAMP/cGMP phosphodiesterases reduces 2′,3′-cGMP level of *N. benthamiana* plants expressing *RBA1*. Empty vector (EV) or constructs indicated were co-expressed with *RBA1* in *N. benthamiana* plants and levels of 2′,3′-cGMP in leaves of the plants were analyzed as described in Figure 5D. The level of 2′,3′-cGMP in *N. benthamiana* expressing EV+EV was normalized to 1.0.

(F) Protein blots for the experiments shown in Figure 6C. Total protein was extracted from 2 dpi leaves shown in Figure 6C and subjected to immunoblot using antibodies indicated. Ponceau staining is shown to indicate loading.

**Figure S7. Model on signaling mechanism induced by 2**′,**3**′**-cAMP/cGMP, related to Figure 7**

(A) Delivery of HopBA1 via Pf0-1 triggers cell death in *Arabidopsis* accession Ag-0 at 48 hpi. Bacteria were injected at OD 0.2 in 10 mM MgCl_2_.

(B) Top: 2′,3′-cAMP/cGMP levels in WT *N. benthamiana* or *epss* quadruple mutant (*eds1, pad4, sag101a, sag101b*) plants expressing *RBA1*. Extract of WT *N. benthamiana* (N.b) or *epss N. benthamiana* (*epss*)was analyzed by LC-MS. Bottom: Protein blots for RBA1 expressed in WT *N. benthamiana* and *epss N. benthamiana*. Total protein was extracted from 2 dpi leaves and subjected to immunoblot using antibodies indicated. Ponceau staining is shown to indicate loading. Cell death was visually assessed and photographed at 5 dpi. The numbers in parentheses indicate the numbers of leaves displaying cell death out of the total number of leaves infiltrated.

(C) L7^TIR^ displays no 2′,3′-cAMP/cGMP synthetase activity when ATP/GTP are used as substrate. 100 μM ATP/GTP or 100 ng PCR product were incubated with L7 ^TIR^ and measured by LC-MS as described in Figure 1E and 1G.

(D) RPP1^TIR^ displays 2′,3′-cAMP/cGMP synthetase activity. The assays were conducted as described in Figure 1E and 1G.

## Experimental Procedures

### Recombinant protein expression and purification

TX0 (At1g57630, residues 1-172) was cloned into pMal C2X vector and expressed in *E. coli* as MBP-fusion proteins with an additional C-terminal GST tag, TX7 (At1g57850, residues 1-192), AbTIR (residues 1-269), L6^TIR^ (residues 26-231) and L7^TIR^ (residues 26-231) were cloned into pMal C2X vector and expressed in *E. coli* as MBP-fusion proteins with additional C-terminal 6 × His tags. The constructs were transformed into the *E. coli* strain BL21 (DE3) (Novagen) at 42°C with 90 s heat shock and the cell cultures were grown at 37°C to OD_600_ of 0.6. Isopropyl-β-D-thiogalactoside (IPTG, Sigma) was added to induce protein expression at 18°C for 16 h. The *E. coli* cells were harvested and lysed by sonification in buffer containing 25 mM Tris-HCl at pH 8.0, 150 mM NaCl, and 15 mM imidazole. The cell lysates were centrifuged at 30,000 *g* for 1.5 h. The supernatant containing soluble proteins was collected and allowed to flow through Ni-NTA resin (GE Healthcare). After washing with three column volumes of sonification buffer, the fusion proteins were eluted in the buffer containing 25 mM Tris-HCl at pH 8.0, 150 mM NaCl, and 150 mM imidazole. For TX0 and TX7, tags could not be completely removed by PreScission protease (GE Healthcare). *Arabidopsis* Nudt7 (residues 1-322), Nudt7^E154Q^ (residues 1-322), *Phytophthora sojae* Avr3b^p6497^ (residues 19-314), *Xanthomonas euvesicatoria* XopQ (residues 85-465), mouse CNPase1 (mCNPase1, residues 161-380) and *Arabidopsis* CPDase (AtCPDase, residues 1-181) were cloned into pET6p-1 vector and expressed in *E. coli* as N-terminal GST-fused proteins. The same protocol as described above was used for *E. coli* culturing. The *E. coli* cells were harvested and lysed by sonification in buffer containing 25 mM Tris-HCl pH 8.0, 150 mM NaCl. The cell lysates were centrifuged at 30,000 *g* for 1.5 hr. The supernatant containing soluble proteins were collected and allowed to flow through GST4B resin (GE Healthcare). After washing with three column volumes of sonification buffer, the fusion proteins were incubated with PreScission at 4°C overnight to remove the N-terminal GST tag and the digested proteins flowed through the columns in the buffer containing 25 mM Tris-HCl pH 8.0, and 150 mM NaCl. WT RBA1 (residues 1-363) and RBA1 mutant proteins were cloned into the pFastBac 1 vector (Invitrogen) with an N-terminal SUMO-6×His tag. The constructs were used for generating recombinant baculovirus in Sf21 insect cells (Invitrogen). RBA1 and RBA1 mutant proteins were expressed in Sf21 insect cells with recombinant baculovirus infection at 28 °C for 48 hr. The infected cells were harvested and lysed by sonification in buffer containing 25 mM Tris-HCl pH 8.0, 150 mM NaCl, and 15 mM imidazole. The cell lysates were centrifuged at 30,000 *g* for 1.5 h. The supernatant containing soluble proteins were collected and allowed to flow through Ni-NTA resin. After washing with three column volumes of sonification buffer, the fusion proteins were incubated with PreScission protease at 4°C overnight to remove the N-terminal SUMO-6×His tag and the digested proteins flowed through the columns in the buffer containing 25 mM Tris-HCl pH 8.0, and 150 mM NaCl.

### *In vitro* NADase assay

Purified WT L7^TIR^ and mutant proteins (20 μM), AbTIR (20 μM), WT RBA1 and mutant proteins (10 μM) were used for NADase assays. Proteins were individually incubated with 100 μM NAD^+^ (final concentration) and 10 mM MgCl_2_ in buffer containing 100 mM NaCl, 25 mM Tris-HCl pH 8.0. The total volume for each reaction was 100 μl. Reactions were performed in a thermoshaker at 25°C for 16 hr. After reaction, samples were centrifuged and immediately applied for high performance liquid chromatography (HPLC) analysis.

HPLC was performed on an Agilent 1260 bioinert HPLC system using a Synergi Fusion-RP 80 Å (4.6 × 150 mm, 4 μm) (Phenomenex) column. The samples were measured via an 8-min method. Samples (10 μl) were injected at 550 μl/min with ammonium formate (5 mM) in water and methanol used as mobile phases A and B, respectively. The elution profile was as follows: 0 to 3 min, 10 to 70% B; 3 to 6 min, 70% B; 6 to 6.1 min, 70 to 10% B; 6.1 to 8 min, 10% B. The autosampler temperature was maintained at 4°C and the column temperature at 25°C. UV signals were detected at 260 nm. Reference standards were used to determine respective retention times. The integrations of peak area were used to calculate relative concentrations.

### Site-directed mutagenesis

*At*NUDT7^E154Q^ construct was introduced using overlap extension PCR. All other mutations of the constructs used for *in vitro* and *in vivo* assays were introduced using a Q5 site-directed mutagenesis kit (NEB). All constructs were verified by DNA sequencing.

### In vitro nuclease activity assay

*Arabidopsis* genomic DNA, *Arabidopsis* total RNA, PCR product and plasmid (100 ng for each) were individually incubated with purified RBA1 and RBA1 mutant proteins (200 nM for each), TX0, TX7, AbTIR, L7^TIR^ and L7^TIR^ mutant proteins (1.0 μM for each) in the buffer containing 25 mM Tris-HCl pH 8.0 and 150 mM NaCl at 25°C for 16 hr. After reaction, the products were mixed with DNA loading buffer and visualized by agarose gel. Reaction products were visualized 6 hr after reaction when RNA was used as substrate.

Nuclease activity was further measured by DNase I Assay Kit (Fluorometric) (ab234056) with a fluorescent DNA probe as the substrate. The purified TIR proteins (5.0 μM) individually were incubated with the fluorescent DNA probe (25 μM) (Abcam). The samples were measured by microplate reader SynergyH1 (BioTek) with excitation/emission 646/686 nm fluorescence.

### Production and detection of 2′,3′-cNMP *in vitro*

The purified RBA1 and RBA1 mutant proteins (5 μM for each), TX0, TX7 AbTIR, L7^TIR^ and L7^TIR^ mutant proteins (1 μM for each) proteins were incubated with 100 ng *Arabidopsis* genomic DNA or *Arabidopsis* total RNA in the buffer containing 25 mM Tris-HCl pH 8.0 and 150 mM NaCl. 2.5μL RNase T1 (Thermo Scientific) or DNase I (Jena Bioscience) was incubated with 100ng *Arabidopsis* total RNA or gDNA. The total volume for each reaction was 100 μl. After incubation at 25°C for 16 hr, the samples were centrifuged at 12,000 g for 10 min, and the supernatant was applied to LC-MS/MS for metabolite identification and quantification.

### Negative staining

For quality examination and overall structure analysis, TX0, RBA1, L6^TIR^, L7^TIR^ and MBP tagged L7^TIR^ were subjected to Superose 6 10/300 gel filtration column (GE Healthcare). Fractions eluted at void volume were collected. 3 μl of each protein (0.1mg/ml) was applied to formvar carbon grids glow-discharged for 45 s at high level in Harrick Plasma after 2 min evacuation. The sample stayed on grid for 1 min at room temperature and dried by touching the edge of a filter paper. 3 μl 2% uranyl acetate was then applied to the grid for 45 s. The excess stain was removed. Grids were dried at room temperature for 2 min. Samples were imaged using Hitachi H-7650 120 kV transmission electron microscope (TEM) and images were recorded with an AMT XR-41 camera.

### Cryo-EM sample preparation and data collection

For structural determination, MBP tagged L7^TIR^ was incubated with by PreScission protease (GE Healthcare) at 4°C or 22°C for 12 hr to remove the MBP tag. The L7^TIR^ protein was further purified by size-exclusion chromatography using Superose6 10/300 gel filtration column in buffer containing10 mM Tris-HCl pH 8.0, 100 mM NaCl. 2.5 μl L7^TIR^ protein (1mg/ml) was applied to holey carbon grids (Quantifoil Au 1.2/1.3, 300 mesh) glow-discharged for 30 s at high level in Harrick Plasma after 2 min evacuation. Grids were then blotted on filter paper (Ted Pella Inc.) for 3 s at 4°C with 100% humidity and plunge-frozen in liquid ethane using FEI Vitrobot Marked IV. Several other conditions were tested with the hope to capture intermediate states, MBP-L7^TIR^ samples were cleaved by PreScission protease at room temperature for 4 hours, then further purified and blotted following similar protocol.

Two datasets of the L7^TIR^ filaments were collected: 4°C processed dataset on a Titan Krios2 electron microscope operated at 300 kV, equipped with Gatan K2 Summit direct electron detector and a Gatan Quantum energy filter, and the 22°C processed dataset on Titan Krios2 with Gatan K3 detector later on.

### Cryo-EM data processing

Processing of the cryo-EM data followed a previous workflow used for reconstructing thin flexible helical complexes (Gong et al., 2021). Firstly, Motion Correction and CTF determination were carried out using corresponding modules in RELION. An in-house picking script designed specifically for filament picking was then used to select end-to-end filament coordinates (https://github.com/Alexu0/Cryo-EM-filament-picker). After segment extraction, coordinates and raw images were imported to cryoSPARC for binned 2D classification. Then selected particle coordinates were transferred back to RELION for helical 2D and 3D classification, 3D helical refinement, and final refinement and masked polishing.

In preliminary non-helical 2D analysis, 150 Å appeared to be the basic repeat unit of the physical rise after rotating the filament by either 180° or 360°. In addition, it appeared that there was a smaller helical unit around 35 Å. Then we exhausted and tested all the potential combinations of helical parameters that fit with these two observations. The preliminary helical parameter hits emerged to be +-43°, and 34 Å. After iterative 3D classification and optimization, it turned out that Initial State Complex had a helical parameter of 159.2°, 16.56 Å (mathematically equivalent to -41.6°, 33.12 Å). Intermediate State Complex was -48.6°, 33.51 Å. End State Complex was -49.3°, 33.43 Å. Based on Gold Standard FSC = 0.143 measurement, the initial complex, intermediate complex and the end state complex maps were refined to 3.4 Å, 3.5 Å and 3.0 Å, respectively.

### Model building

Previous TIR domain crystal structure was used as initial model. Model refinement was achieved by fitting PDB 6O0W into a single TIR domain density of the end state complex. Phenix real space refinement module was used to optimize the model fitting. Final model to density FSC (0.143) = 2.95 Å, while mask to density CC values (CC-mask, volume, peak, box) were within the range of 0.718 to 0.791. An extra blob of density was identified inside the nucleotide binding pocket. Extensive small molecule docking identified 2’,3’-cyclic AMP as the best fit to accommodate this extra density.16 refined L7^TIR^ domain models were docked into initial state complex, and the remaining dsDNA like density was fitted with 43 bp A-form dsDNA, model generated by Chimera. For every 11bp DNA, there was 2-3 base pairs found to be stretched, and possessing high slide and twist values, after Phenix real space optimization of the model fitting. This 43bp-16 TIR model was used to analyze how TIR domains oligomerize and cooperatively recognize nucleic acids. From several intermediate state complex structures, one representative density which showed all double stranded region hydrolyzed down to 3 remaining base pair was selected for model building. Using Coot and Chimera, nucleotides were manually added in to best represent the density. A detailed 4 L7^TIR^ domain model with the bound nucleic acids was built, with the aim of analyzing the nucleic acid-TIR interface during the progressive hydrolysis.

Two L7^TIR^ domain models together with the bound single stranded DNA was docked into the end state complex map to determine the complex conformation at the end of nucleic acid hydrolysis. Nucleotides were manually docked in, using Coot and Chimera, later optimized using Phenix real space refinement. This model highlighted additional TIR-TIR interactions and how the last remain single stranded nucleic acids remained bound. At this conformation, the 2’3’-cNMP density inside the TIR domain was most obvious among all complexes.

### Protein expression in *N. benthamiana* and western blot analysis

For *N. benthamiana* transient expression, *RBA1* (residues 1-363), *AtNUDT7* (residues 1-322), *PDE9* (residues 1-533), *mCNPase1* (residues 161-380), *AtCPDase* (residues 1-181), *Avr3b* (residues 19-314), were cloned into the pENTR/D-TOPO vector (Thermo Fisher Scientific, K240020). The obtained pENTR/D-TOPO *RBA1* and *RBA1* mutant plasmids were LR-recombined into pXCSG vectors with a C-terminal MYC tag, the obtained pENTR/D-TOPO *AtNUDT7* and *PDE9* plasmids were LR-recombined into pEGAD vectors with a C-terminal GFP tag and the obtained pENTR/D-TOPO *mCNPase1, AtCPDase, Avr3b* plasmids were LR-recombined into pXCSG vectors with a C-terminal 3×HA tag. All the constructs were verified by DNA sequencing. Generated binary constructs were transformed into *Agrobacterium tumefaciens* (Rhizobium radiobacter) GV3101 pMP90RK via electroporation.

*RBA1* and *RBA1* mutants were expressed in leaves of four-week-old *N. benthamiana* plants using *agrobacteria*-mediated transient expression assays in the presence of the P19 suppressor of RNAi silencing. The final OD600 for each strain was set to 0.8. For co-infiltration, *agrobacteria* carrying TIR and hydrolases expression constructs were mixed at 1:1 ratio. TIRs mixed with the empty vector (EV) were applied as a negative control to rule out the possibility that high titer of *agrobacteria* would affect infection efficiency. To detect RBA1, RBA1 mutants and the co-infiltrated AtNUDT7, AtNUDT7^E154Q^ and PDE9 protein expression, *N. benthamiana* leaves were collected at 2 dpi, snap frozen in liquid nitrogen, and ground to fine powder. The powder (∼100 μl in a tube) was resuspended in 100 μl of urea-SDS sample buffer [50 mM Tris-HCl pH 6.8, 2% SDS, 8 M urea, 2% β-mercaptoethanol, 5% glycerol, protease inhibitor cocktail (Roche), and 0.004% Bromophenol Blue] and vortexed for 10 min at room temperature. No boiling step was included. After centrifugation at 16,000 g for 10 min, 10 μl of the supernatant was loaded onto 8% SDS-PAGE and proteins were blotted onto a PVDF membrane.

To detect co-infiltrated mCNPase, AtCPDase and Avr3b proteins expression, 300 mg *N. benthamiana* leaves were collected at 2 dpi and snap frozen in liquid nitrogen. All further steps were performed at 4°C if not mentioned otherwise. Tissue samples were ground to fine powder and resuspended in 1 ml of IP buffer [100mM Tris-HCl pH 7.5, 150 mM NaCl, 5mM EDTA, 2mM EGTA, 5% (v/v) glycerol, 1% Triton, 2% (w/v) PVPP, 10mM Dithiothreitol (DTT) 2% (v/v), protease inhibitor cocktail (Roche), 0.5 mM PMSF] and incubated for 1 h with gentle rotation. Debris was removed by centrifugation at 21,000 g for 15 min. Immunoprecipitations were performed with 5 µl rabbit polyclonal anti-HA antibody (Abcam; 9110) coupled to 50µl bed volume Protein A agarose (Roche) per sample. After 4 h of incubation under constant rotation, beads were washed 5 times in 1 mL IP buffer (without PVPP) and eluted by boiling in 100 μL of 2× Laemmli sample buffer for 10 min at 95°C. 10 μl of the was loaded onto 10% SDS-PAGE and proteins were blotted onto a PVDF membrane.

Immunoblot assays were performed using monoclonal rat anti-HA antibody (Sigma Aldrich, 11867423001), mouse anti-Myc antibody (ThermoScientific R950-25), and rabbit anti-GFP antibody (Chromtek.pabg1-100) diluted 1:3000 in 1X TBS, 0.1% Tween-20 with 5% non-fat milk powder. Proteins were detected with a Gel Doc XR+ Gel Documentation System by using ECL SuperSignal West Femto Maximum Sensitivity Substrate and ECL Western Blotting substrate (Thermo Scientific) in a ratio of 2:1.

### Cell death quantification in *N. benthamiana*

*RBA1* and *RBA1* mutants and combinations with hydrolase protein *NUDT7, PDE9, mCNPase, AtCPDase* and *Avr3b* were transiently expressed in *N. benthamiana* as described above and three individual *agrobacteria*-infiltrated leaf zones were used for ion leakage assays. Five 8-mm leaf discs from *N. benthamiana* agroinfiltrated leaves were taken at 3 dpi, washed in 10 to 20 mL of milliQ water (18.2 MΩ*cm, mQ) for 30 to 60 min, transferred to a 24-well plate with 1 mL mQ in each well, and incubated at room temperature. Ion leakage was measured at 0 and 6 h with a conductometer Horiba Twin Model B-173. Statistical analysis was performed on conductivity data via Tukey′s HSD (honestly significant difference) test (a=0.05). Images of *agrobacteria*-infiltrated leaf spots were taken at 4 to 5 dpi.

### HopBA delivery assay

Cloning of the HopBA1 effector into bacterial type III secretion vector pCK014 to mobilize into the Pseudomonas fluorescens strain Pf0-1was performed as described (Gantner et al., 2018). Overnight grown Pf0-1 carrying HopBA-1 or empty vector bacteria were resuspended in 10 mM MgCl_2_ to OD600 of 0.2 and infiltrated with a needleless syringe into rosette leaves of 5-6 weeks old Arabidopsis Ag-0 plants before noon. 3 infiltrated leaves were harvested from each pot, leaves from 3 pots were harvested and snap-frozen at 24 or 36 hpi for 2′,3′-cAMP/cGMP detection. Images of infiltrated leaves were taken at 48 hpi.

### *In vivo* 2′,3′-cNMP extraction and detection

2′,3′-cNMP were extracted from *N. benthamiana* or Arabidopsis as described (Van Damme et al., 2014). For WT and *RBA1* mutants infiltrated or *RBA1*-hydrolase co-infiltrated *N. benthamina* leaves, 3 individual infiltrated zones were harvested and snap-frozen at 3 dpi. For 4 weeks old Col-0 and *Atnudt7 eds1* Arabidopsis, 3 leaves were harvested from each pot; for 4 weeks old *Atnudt7* leaves, due to the dwarf phenotype, 2 leaves were harvested from each pot. Leaves from 4 pots were harvested and snap-frozen. Frozen plant leaves were homogenized manually with a precooled mortar and pestle with liquid nitrogen. 100 (±5) mg ground tissue was transferred to precooled 2 mL Eppendorf tubes. 600 μl 4% acetic acid with 20 μl 1mM phosphodiesterase inhibitor 3-isobutyl-1-methylxanthine (IBMX, Sigma) and 0.6 μl 1mM spike control 8-Br-2′,3′-cAMP (Biolog) were added to per 100 mg sample. The samples were vortexed for 2 min and subsequently centrifuged for 9 min at 9000 g. The supernatant was transferred into new precooled tubes. 800 μl chloroform was added to each tube. The tubes were gently inverted 6 times and then centrifuged at 6000 g for 6 min. Aqueous phase were collected and extracted again by chloroform. The samples were dried at 30°C in speed vacuum concentrator and reconstituted in 50 μL of water and vortexed for 1 min. The samples were centrifuged 12,000 g for 10 min, and the supernatant was applied to the LC-MS/MS for metabolite identification and quantification.

### Metabolite measurement by LC-MS/MS

Chromatography was performed on a Nexera XR 40 series HPLC (Shimadzu) using a Synergi 4µM Fusion-RP 80 Å 150×2mm column (Phenomenex). The column temperature was maintained at 40°C and the sample tray at 4°C. Samples (10μl) were injected at a flow rate of 0.2 ml/min using 10 mM ammonium formate at pH 4.2 and methanol as mobile phases A and B, respectively. Metabolites were eluted using the profile 0-8 min, 8-90% B; 8-10 min, 90% B; 10-10.1 min 90-8% B; 10.1-20 min, 8% B. The LCMS-8060 triple quadrupole mass spectrometer with electro spray ionization (Shimadzu) was operated in positive mode. Scheduled multiple reaction monitoring (MRM) was used to monitor analyte parent ion to product ion formation. MRM conditions were optimized using authentic standard chemicals including: 2′,3′-cAMP ([M+H]^+^ 330.00>136.20, 330.00>99.15, 330.00>119.15), 2′,3′-cGMP ([M+H]^+^ 346.00>152.15, 346.00 >135.20, 346.00>110.15), 2′,3′-cCMP ([M+H]^+^ 306.00>112.15, 306.00>95.20, 306.00>178.20), 2′,3′-cUMP ([M+H]^+^ 307.00>113.15, 307.00>195.10, 307.00>136.15), 8-Br-2′,3′-cAMP ([M+H]^+^ 407.90 >214.00, 407.90>99.20, 407.90>69.25), 3′,5′-cAMP ([M+H]^+^ 329.80>135.95, 329.80>311.90, 329.80>96.95), 3′,5′-cGMP ([M+H]^+^ 345.90>151.95, 345.90>135.05, 345.90>110.20), 3′,5′-cCMP ([M+H]^+^ 306.00>112.20, 306.00>95.20, 306.00>69.20), 3′,5′-cUMP ([M+H]^+^ 307.00>97.15, 307.00>148.15, 307.00>113.15). Both Q1 and Q3 quadrupoles were maintained in unit resolution. LabSolutions LCMS v5.97 software was used for data acquisition and LabSolutions Postrun for processing (both Shimadzu). Metabolites were quantified by scheduled MRM peak integration using external calibration curves of standard chemicals.

