## Supplemental Figures for "TIR domains of plant immune receptors are 2′,3′-cAMP/cGMP synthetases mediating cell death"

### Figure s1

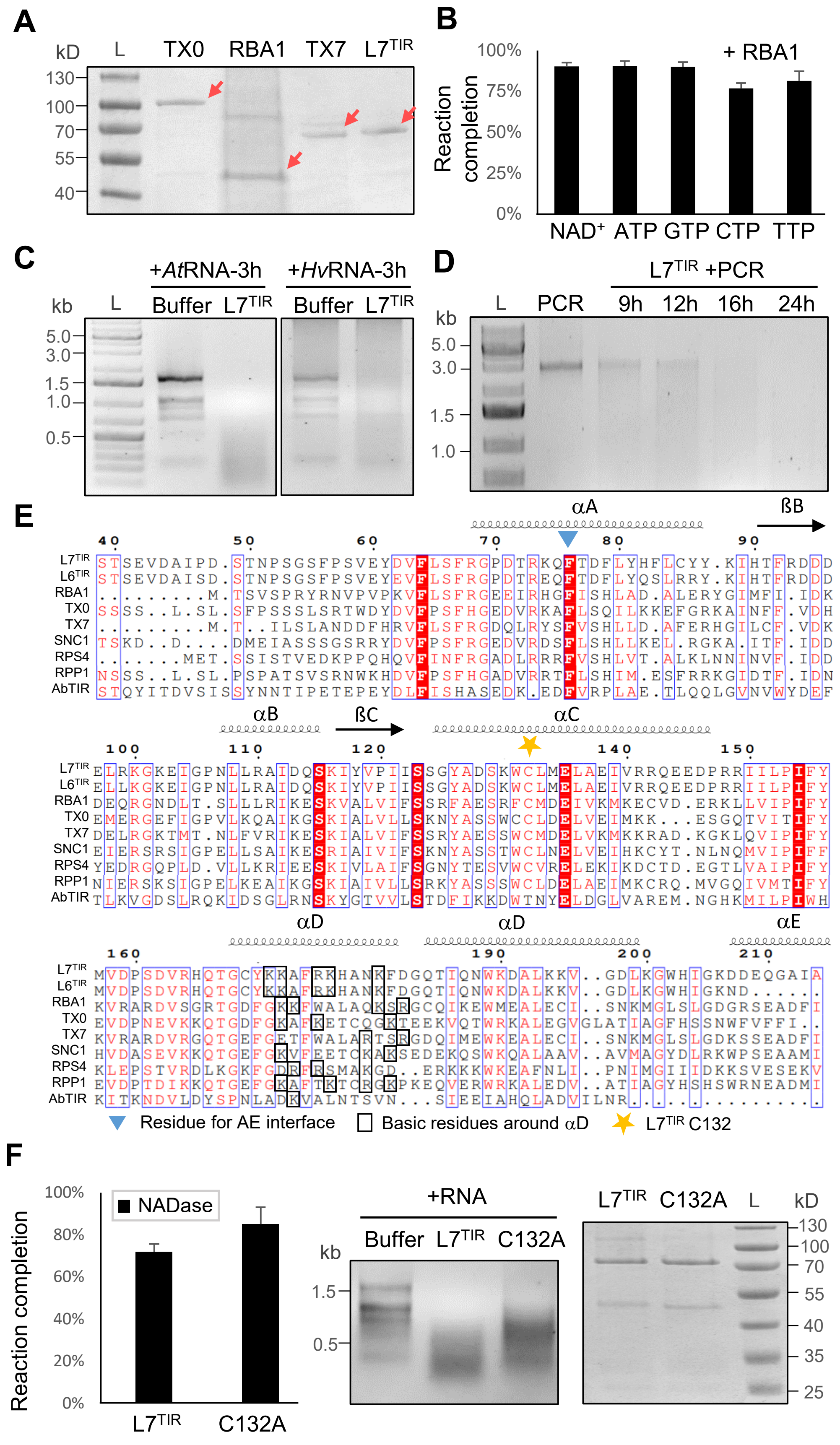
A
kD
L
TX0
RBA1
TX7
L7TIR
130
100
70
55
40
B
100%
75%
50%
25%
0%
Reaction completion
+ RBA1
NAD+
ATP
GTP
CTP
TTP
C
+AtRNA-3h
+HvRNA-3h
kb
L
Buffer
L7TIR
Buffer
L7TIR
5.0
3.0
1.5
1.0
0.5
D
L7TIR +PCR
kb
L
PCR
9h
12h
16h
24h
5.0
3.0
1.5
1.0
E
αA
ßB
40 50 60 70 80 90
L7TIR STSEVDAIPD.STNPSGSFPSVEYDVFLSFRGPDTRKQFTDFLYHFLCYY.KIHTFRDDD
L6TIR STSEVDAISD.STNPSGSFPSVEYEVFLSFRGPDTREQFTDFLYQSLRRY.KIHTFRDDD
RBA1 .........M.TSVSPRYRNVPVPKVFLSFRGEEIRHGFISHLAD.ALERYGIMFI.IDK
TX0 SSSS..L.SL.SFPSSSLSRTWDYDVFPSFHGEDVRKAFLSQILKKEFGRKAINFF.VDH
TX7 .........M.T..ILSLANDDFHRVFLSFRGDQLRYSFVSHLLD.AFERHGILCF.VDK
SNC1 TSKD..D.....DMEIASSSGSRRYDVFPSFRGEDVRDSFLSHLLKEL.RGKA.ITF.IDD
RPS4 ......MET.SSISTVEDKPPQHQVFINFRGADLRRRFVSHLVT.ALKLNNINVF.IDD
RPP1 NSSS..L.SL.PSPATSVSRNWKHDVFPSFHGADVRRTFLSHIM.ESFRRKGIDTF.IDN
AbTIR STQYITDVSISSYNNTIPETEPEYDLFISHASEDK.EDFVRPLAE.TLQQLGVNVWYDEF
αB
ßC
αC
100 110 120 140 150
L7TIR ELRKGKEIGPNLLRAIDQSKIYVPIISSGYADSKWCLMELAEIVRRQEEDPRRIILPIFY
L6TIR ELLKGKEIGPNLLRAIDQSKIYVPIISSGYADSKWCLMELAEIVRRQEEDPRRIILPIFY
RBA1 DEQRGNDLT.SLLLRIKESKVALVIFSSRFAESRFCMDEIVKMKECVD.ERKLLVIPIFY
TX0 EMERGEFIGPVLKQAIKGSKIALVLLSKNYASSWCCLDELVEIMKK...ESGQTVITIFY
TX7 DELRGKTMT.NLFVRIKESKIALVIFSSRYAESSWCMDELVKMKKRAD.KGKLQVIPIFY
SNC1 EIERSRSIGPELLSAIKESRIAIVIFSKNYASSTWCLNELVEIHKCYT.NLNQMVIPIFF
RPS4 YEDRGQPLD.VLLKRIEESKIVLAIFSGNYTESVWCVRELEKIKDCTD.EGTLVAIPIFY
RPP1 NIERSKSIGPELKEAIKGSKIAIVLLSRKYASSSWCLDELAEIMKCRQ.MVGQIVMTIFY
AbTIR TLKVGDSLRQKIDSGLRNSKYGTVVLSTDFIKKDWTNYELDGLVAREM.NGHKMILPIWH
αD
αD
αE
160 190 200 210
L7TIR MVDPSDVRHQTGCYKKAFRKHANKFDGQTIQNWKDALKKV..GDLKGWHIGKDDEQGAIA
L6TIR MVDPSDVRHQTGCYKKAFRKHANKFDGQTIQNWKDALKKV..GDLKGWHIGKND......
RBA1 KVRARDVSGRTGDFGKKFWALAQKSRGCQIKEWMEALECI..SNKMGLSLGDGRSEADFI
TX0 EVDPNEVKKQTGDFGKAFKETCQGKTEEKVQTWRKALEGVGLATIAGFHSSNWFVFFI..
TX7 KVRARDVRGQTGEFGETFWALARTSRGDQIMEWKEALECI..SNKMGLSLGDKSSEADFI
SNC1 HVDASEVKKQTGEFGKVFEETCKAKSEDEKQSWKQALAAV..AVMAGYDLRKWPSEAAMI
RPS4 KLEPSTVRDLKGKFGDRFRSMAKGD..ERKKKWKEAFNLI..PNIMGIIIDKKSVESEKV
RPP1 EVDPTDIKKQTGEFGKAFTKTCRGKPKEQVERWRKALEDV..ATIAGYHSHSWRNEADMI
AbTIR KITKNDVLDYSPNLADKVALNTSVN...SIEEIAHQLADVILNR.............
Residue for AE interface
Basic residues around αD
L7TIR C132
F
100%
80%
60%
40%
20%
0%
Reaction completion
NADase
L7TIR
C132A
+RNA
kb
Buffer
L7TIR
C132A
1.5
0.5
L7TIR
C132A
L
kD
130
100
70
55
40
35
25

### Figure s1 (continued)

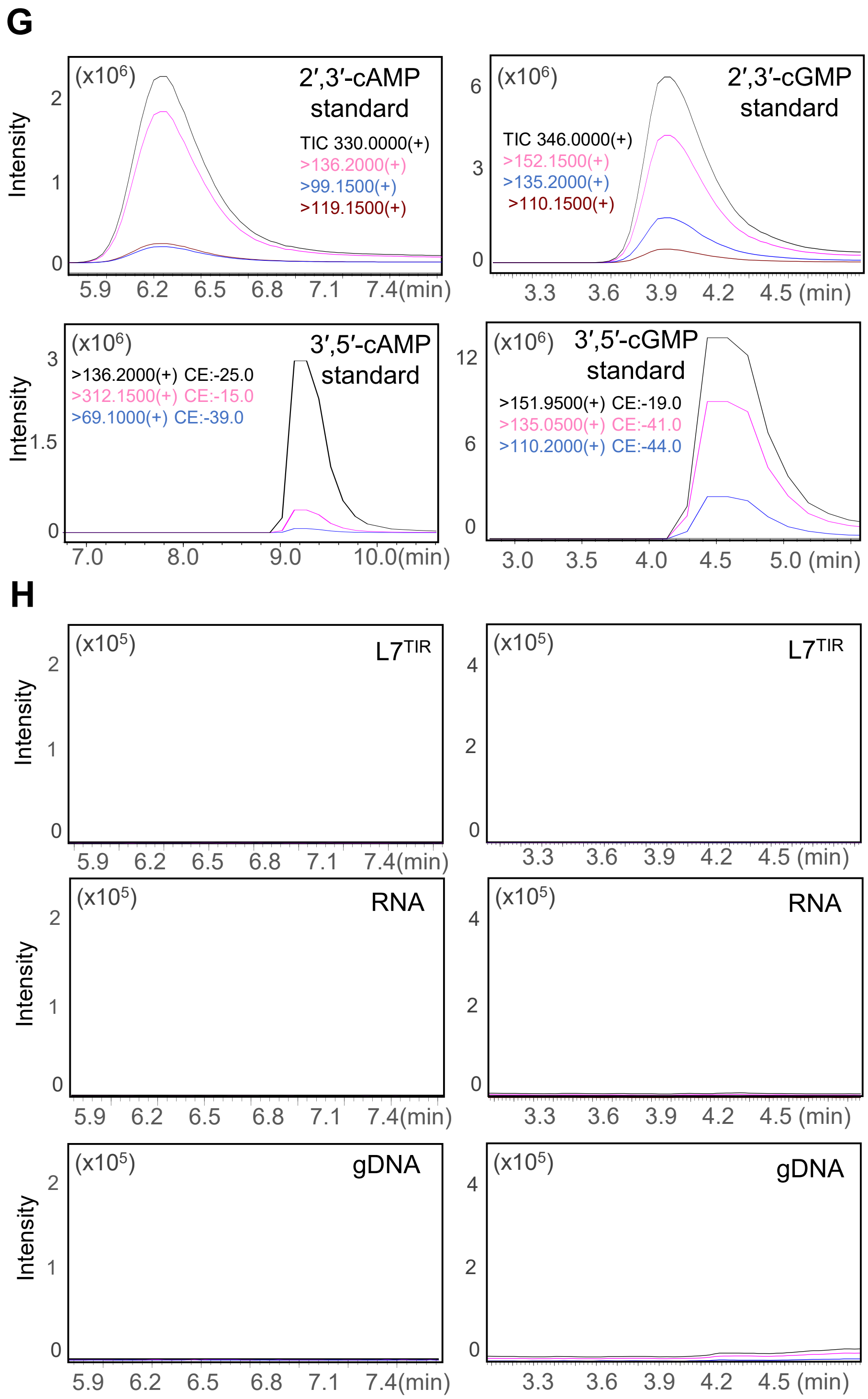

### Figure s1 (continued)

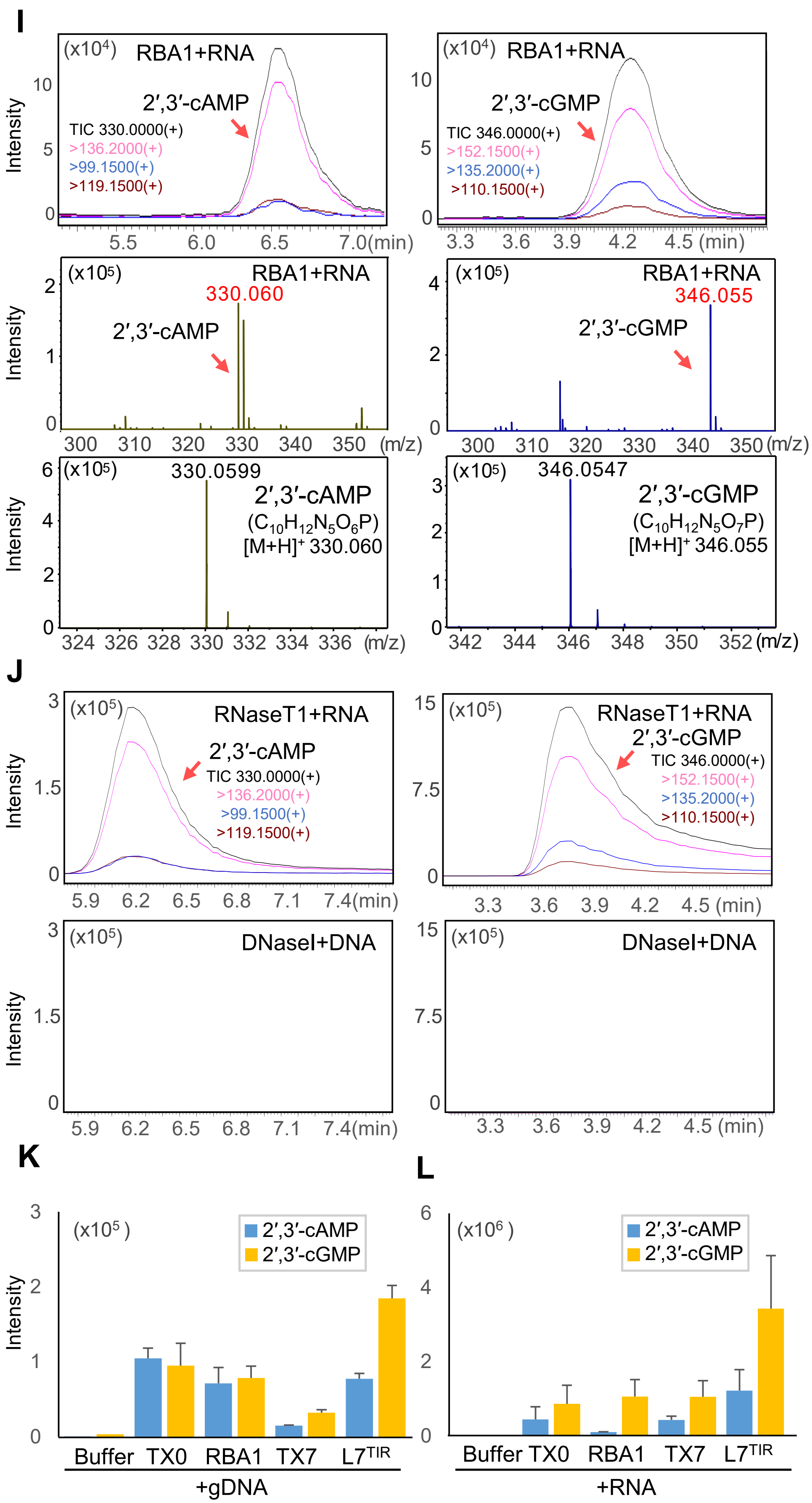

### Figure s2

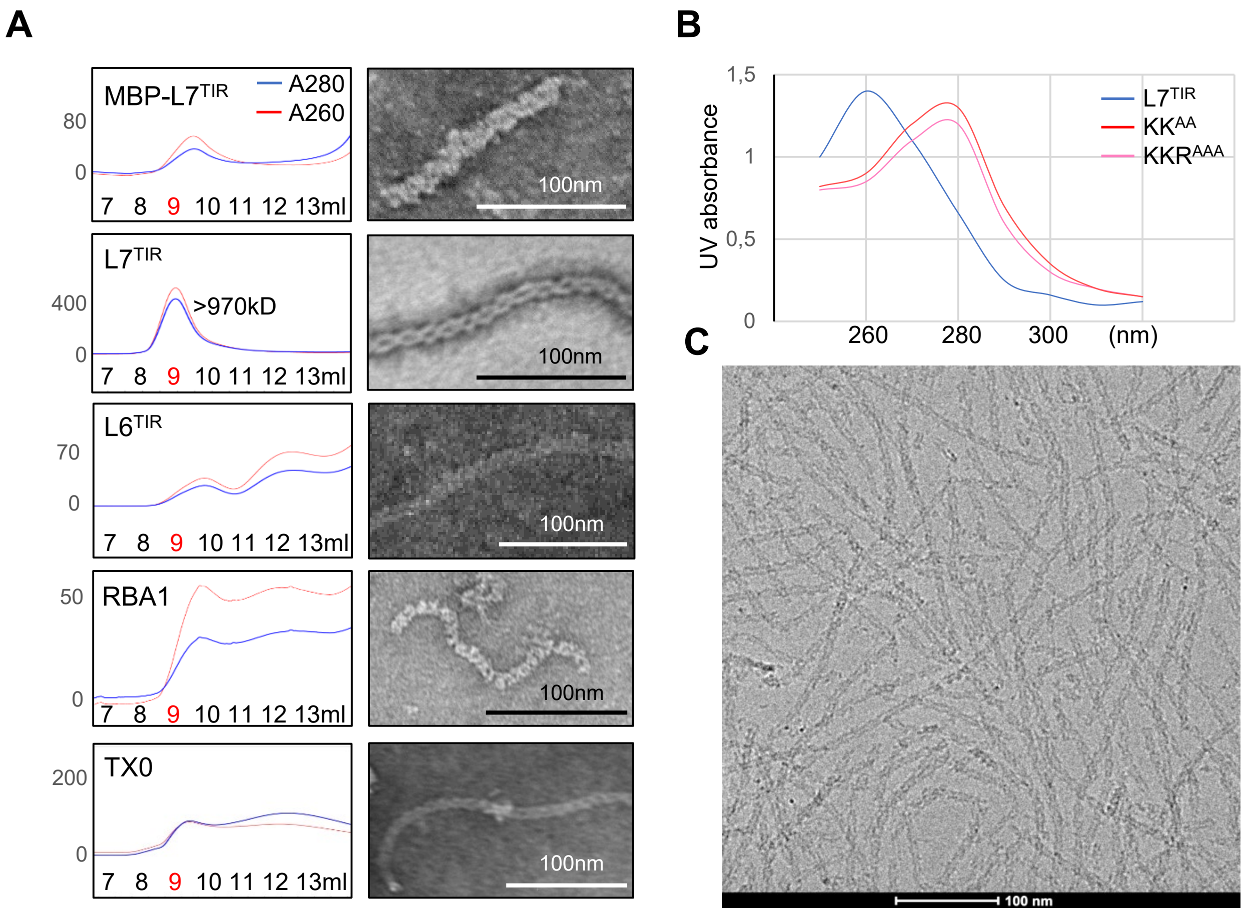
A
MBP-L7TIR
A280
A260
80
0
7 8 9 10 11 12 13ml
100nm
L7TIR
400
>970kD
0
7 8 9 10 11 12 13ml
100nm
L6TIR
70
0
7 8 9 10 11 12 13ml
100nm
RBA1
50
0
7 8 9 10 11 12 13ml
100nm
TX0
200
0
7 8 9 10 11 12 13ml
100nm
B
1,5
1
0,5
0
UV absorbance
260 280 300 (nm)
L7TIR
KKAA
KKRAAA
C
100 nm

**Figure s2 (continued)**

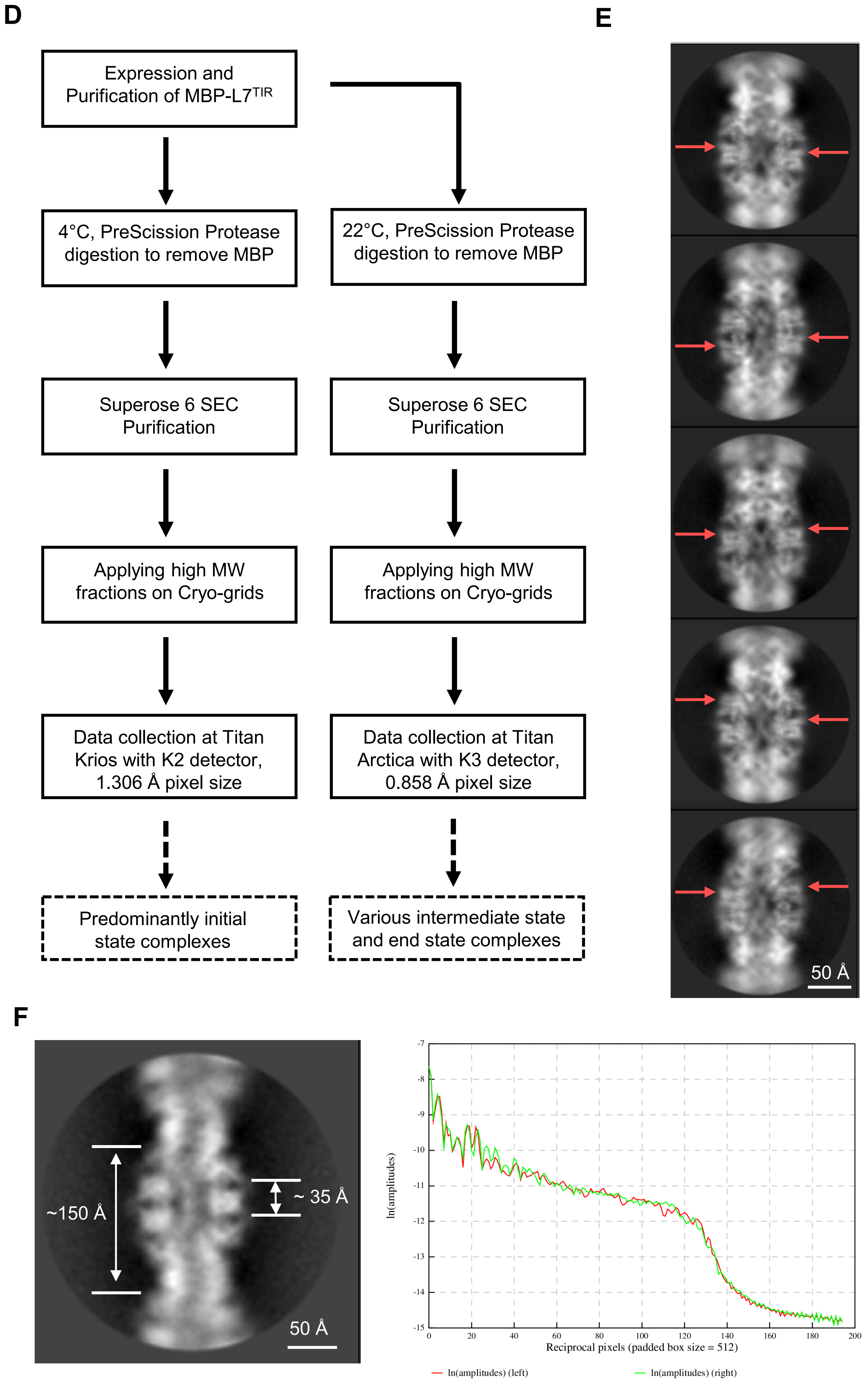

### Figure s2 (continued)

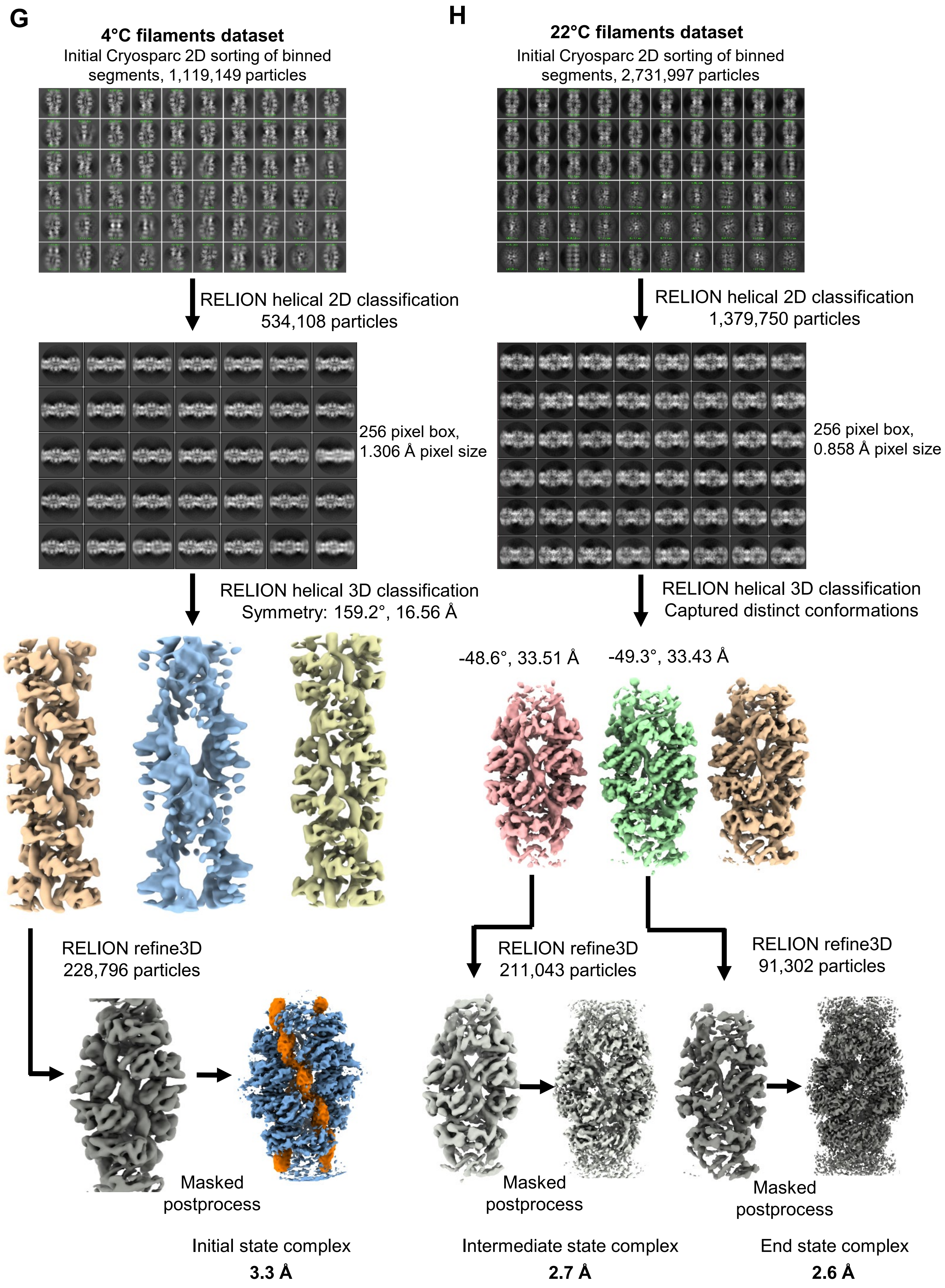

### Figure s2 (continued)

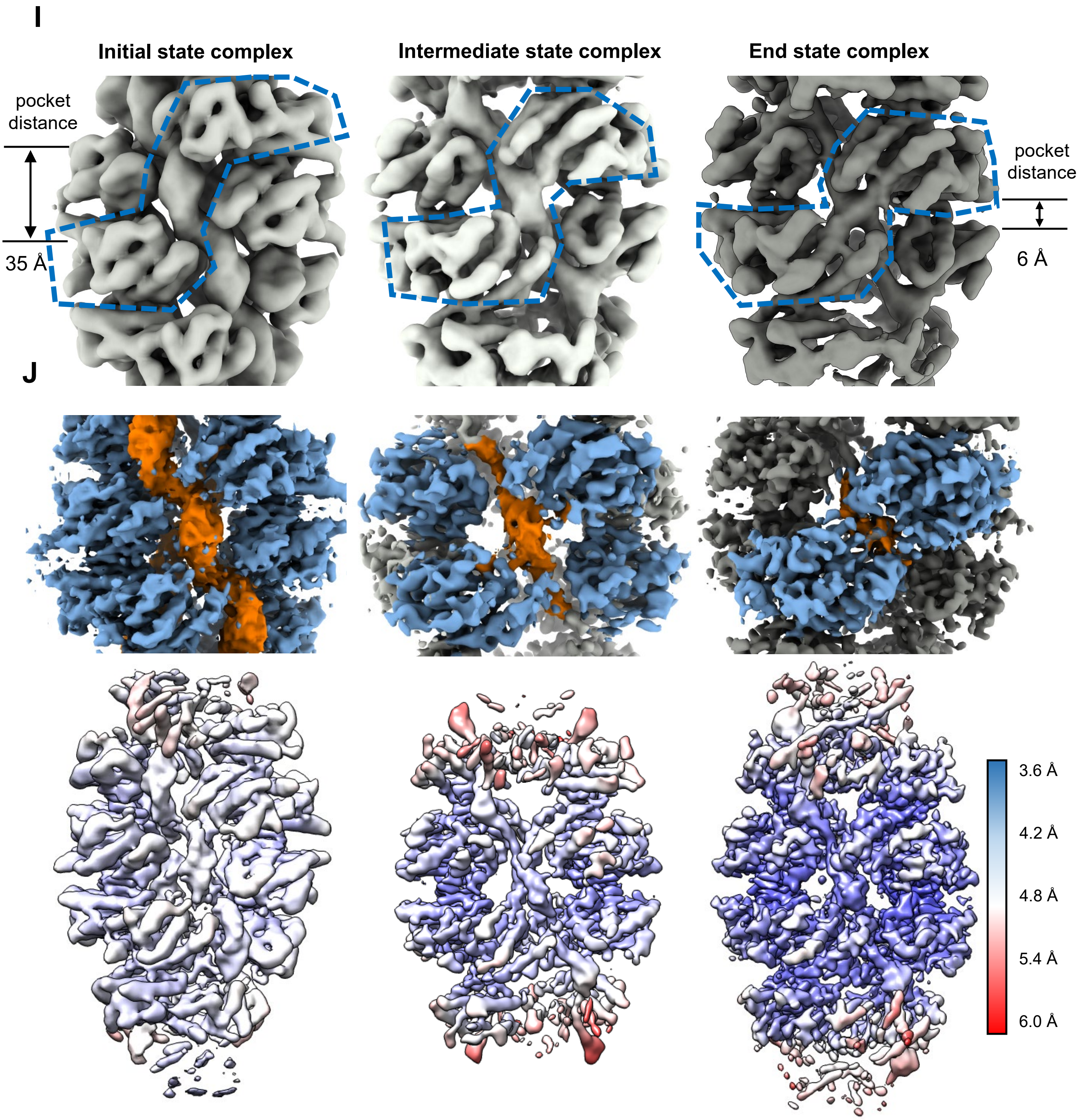

### Figure s2 (continued)

**K**

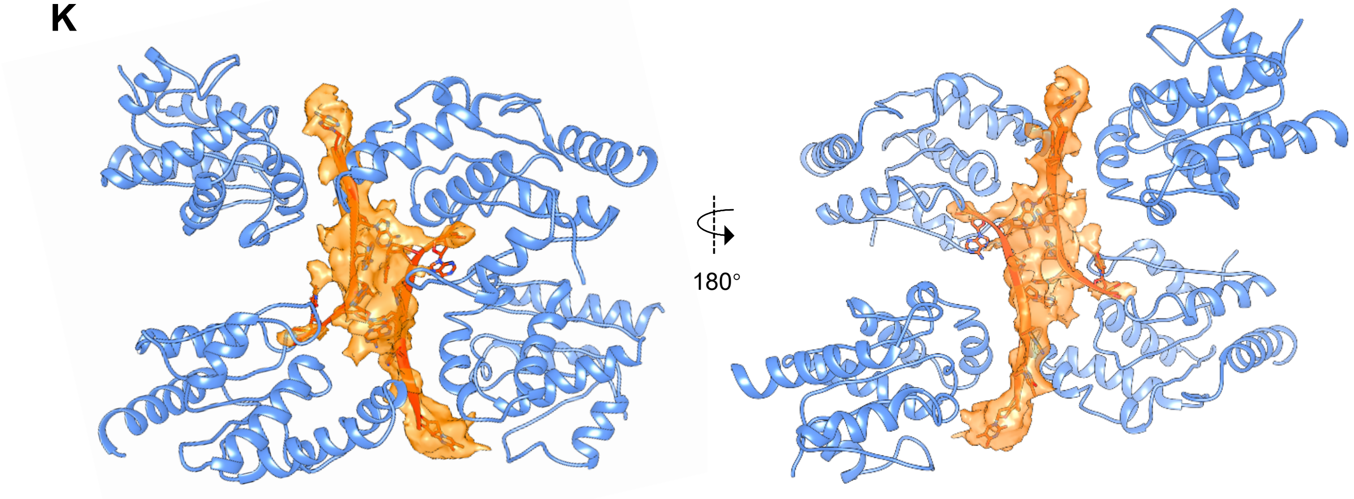


**L**

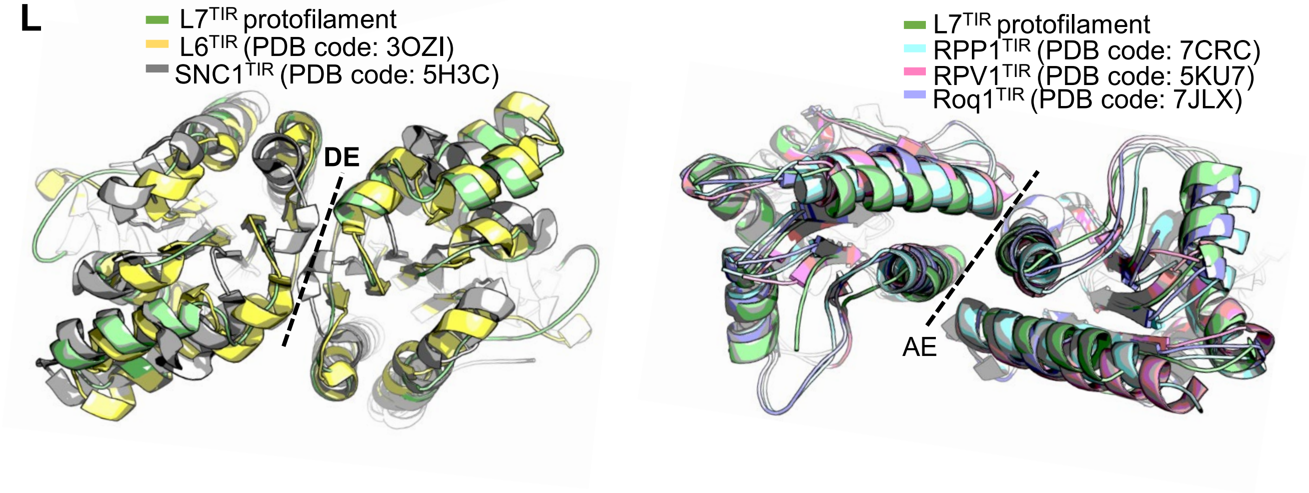

### Figure s2 (continued)

**M**

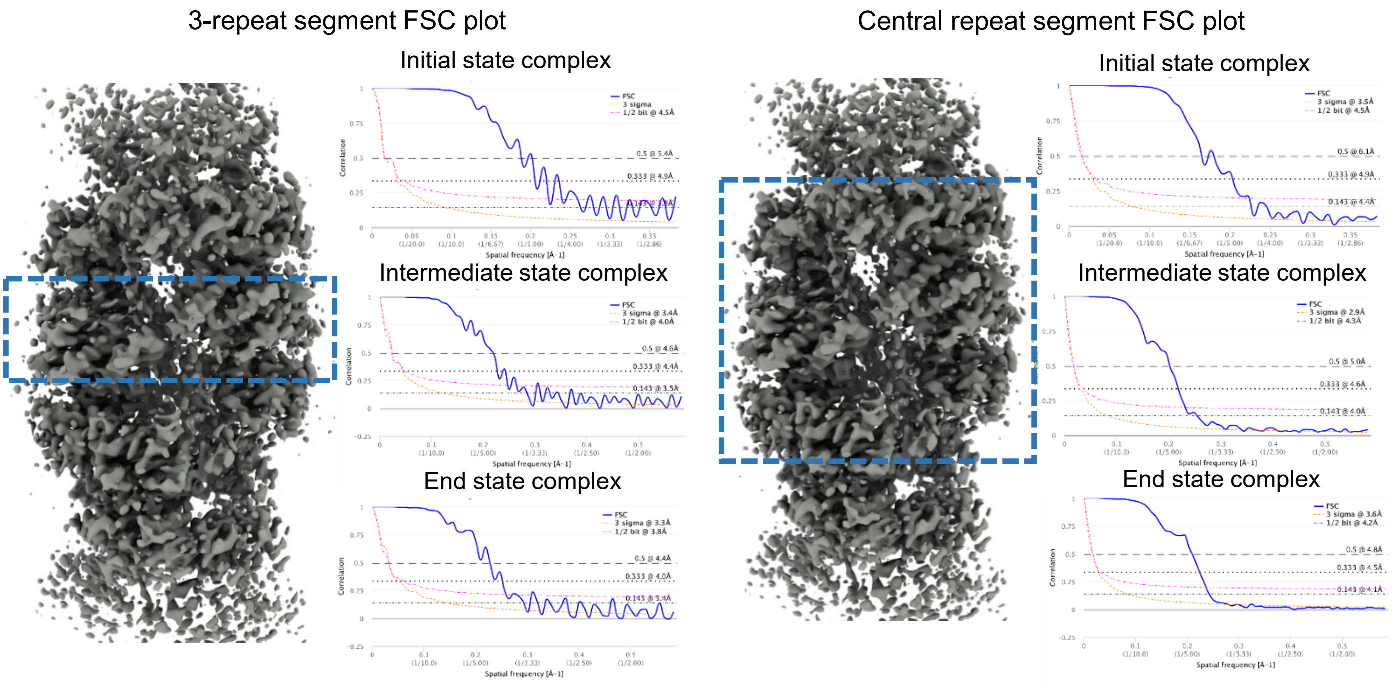


**N**

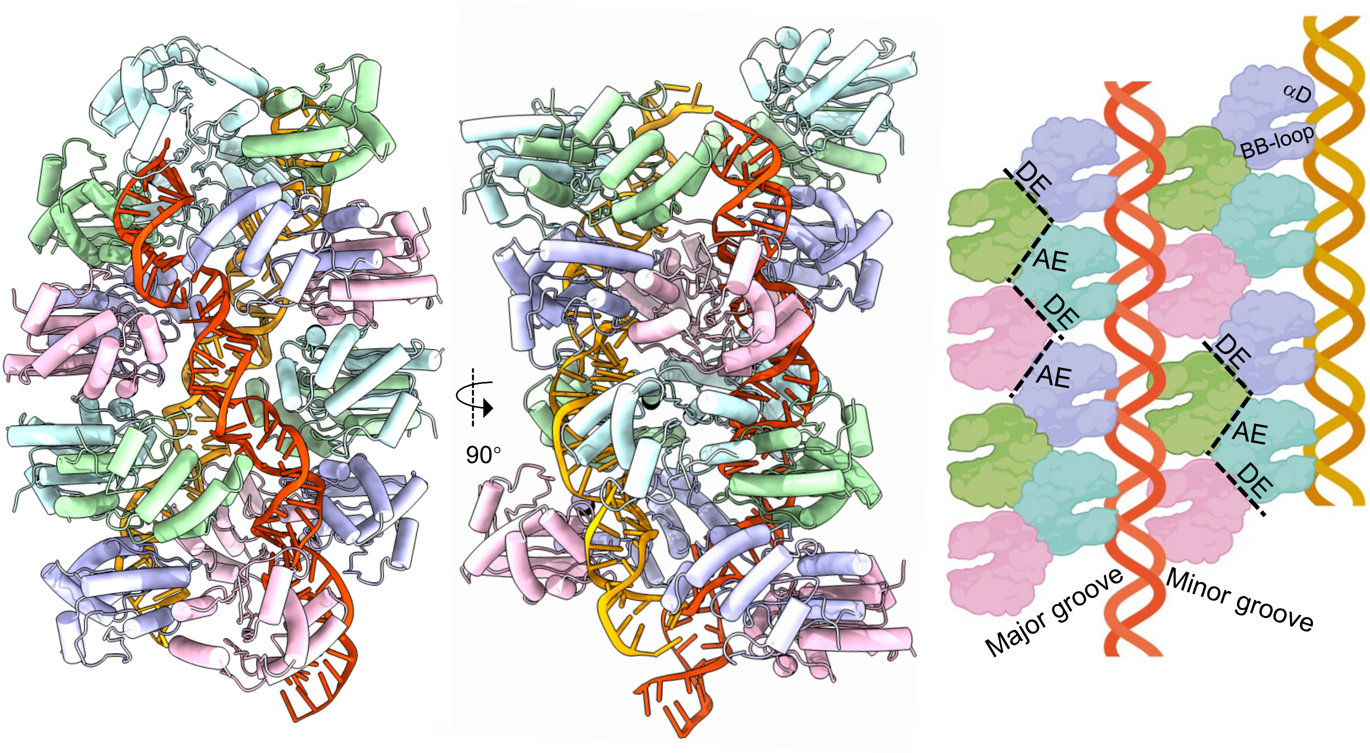

### Figure s3

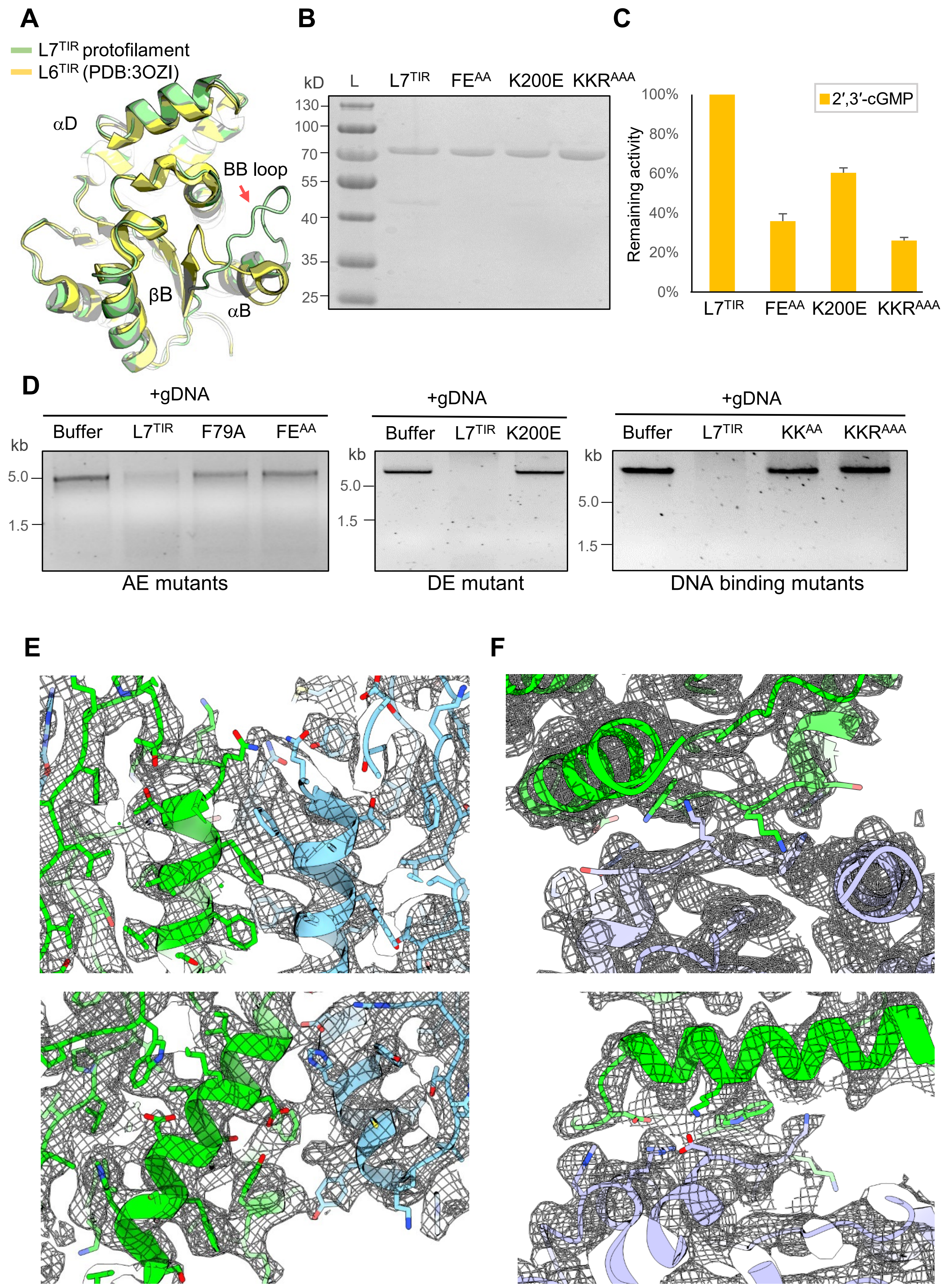
A
L7TIR protofilament
L6TIR (PDB:3OZI)
αD
BB loop
βB
αB
B
kD
L
L7TIR
FEAA
K200E
KKRAAA
130
100
70
55
40
35
25
C
100%
80%
60%
40%
20%
0%
Remaining activity
2′,3′-cGMP
L7TIR
FEAA
K200E
KKRAAA
D
+gDNA
Buffer
L7TIR
F79A
FEAA
kb
5.0
1.5
AE mutants
+gDNA
Buffer
L7TIR
K200E
kb
5.0
1.5
DE mutant
+gDNA
Buffer
L7TIR
KKAA
KKRAAA
kb
5.0
1.5
DNA binding mutants
E
F

### Figure s4

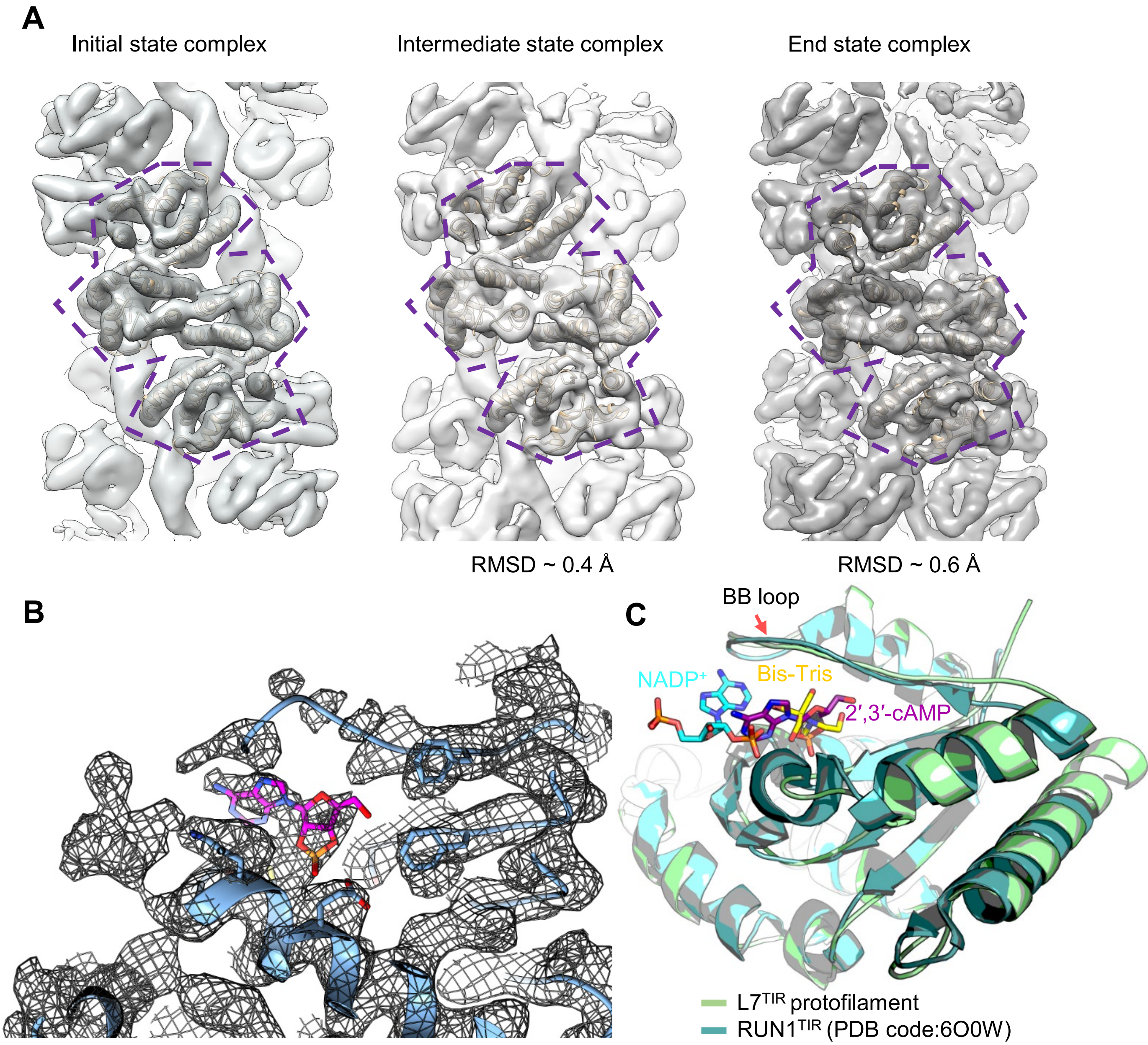
A
Initial state complex
Intermediate state complex
End state complex
RMSD ~ 0.4 Å
RMSD ~ 0.6 Å
B
C
BB loop
NADP+
Bis-Tris
2′,3′-cAMP
L7TIR protofilament
RUN1TIR (PDB code:6O0W)

### Figure s5

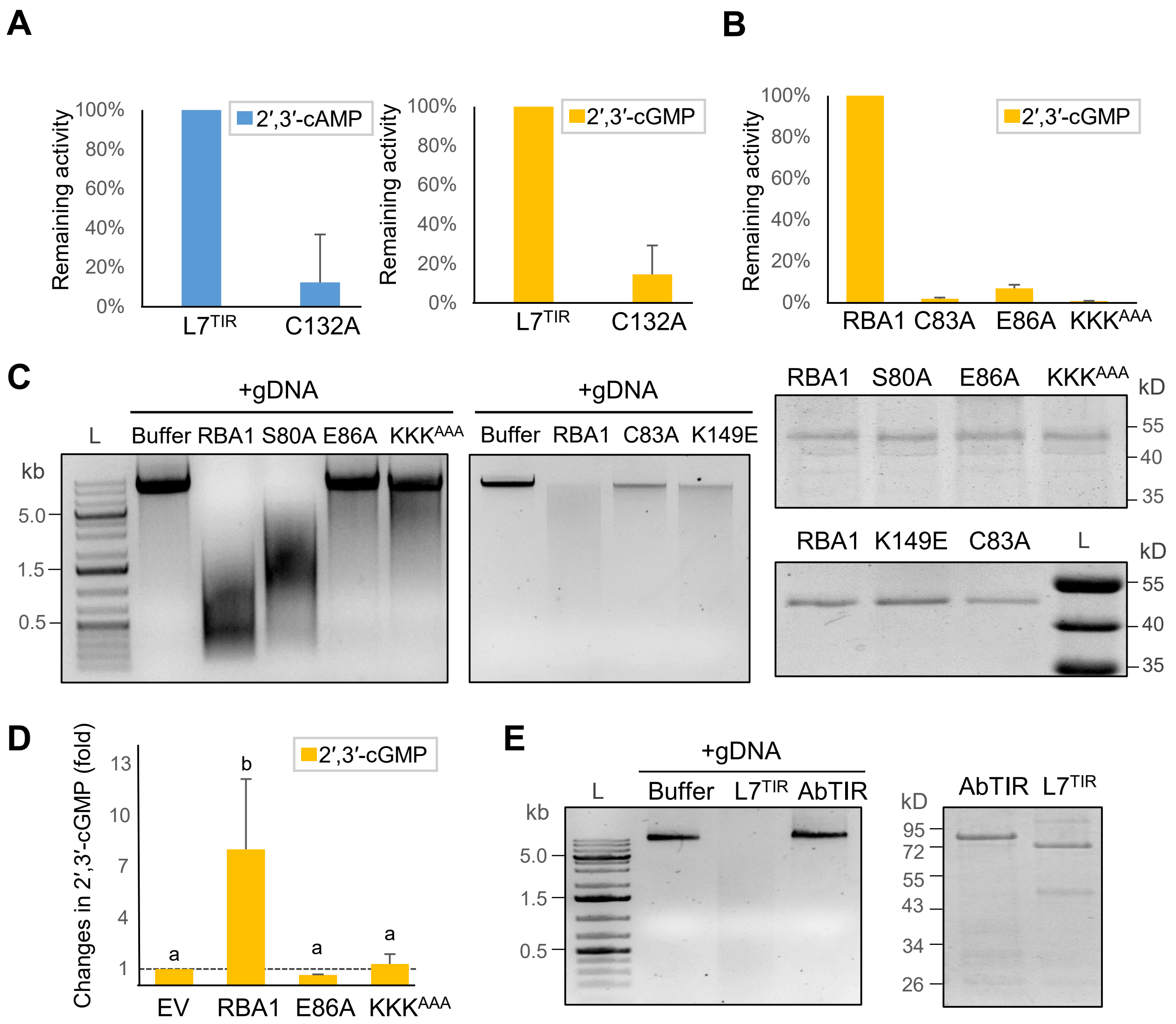

A
Remaining activity
100%
80%
60%
40%
20%
0%
2′,3′-cAMP
L7TIR
C132A
2′,3′-cGMP
B
RBA1
C83A
E86A
KKKAAA
C
+gDNA
L
Buffer
RBA1
S80A
E86A
KKKAAA
kb
5.0
1.5
0.5
+gDNA
Buffer
RBA1
C83A
K149E
RBA1
S80A
E86A
KKKAAA
kD
55
40
35
RBA1
K149E
C83A
L
kD
55
40
35
D
Changes in 2′,3′-cGMP (fold)
13
10
7
4
1
2′,3′-cGMP
a
b
a
a
EV
RBA1
E86A
KKKAAA
E
+gDNA
L
Buffer
L7TIR
AbTIR
kb
5.0
1.5
0.5
kD
95
72
55
43
34
26
AbTIR
L7TIR

### Figure s6

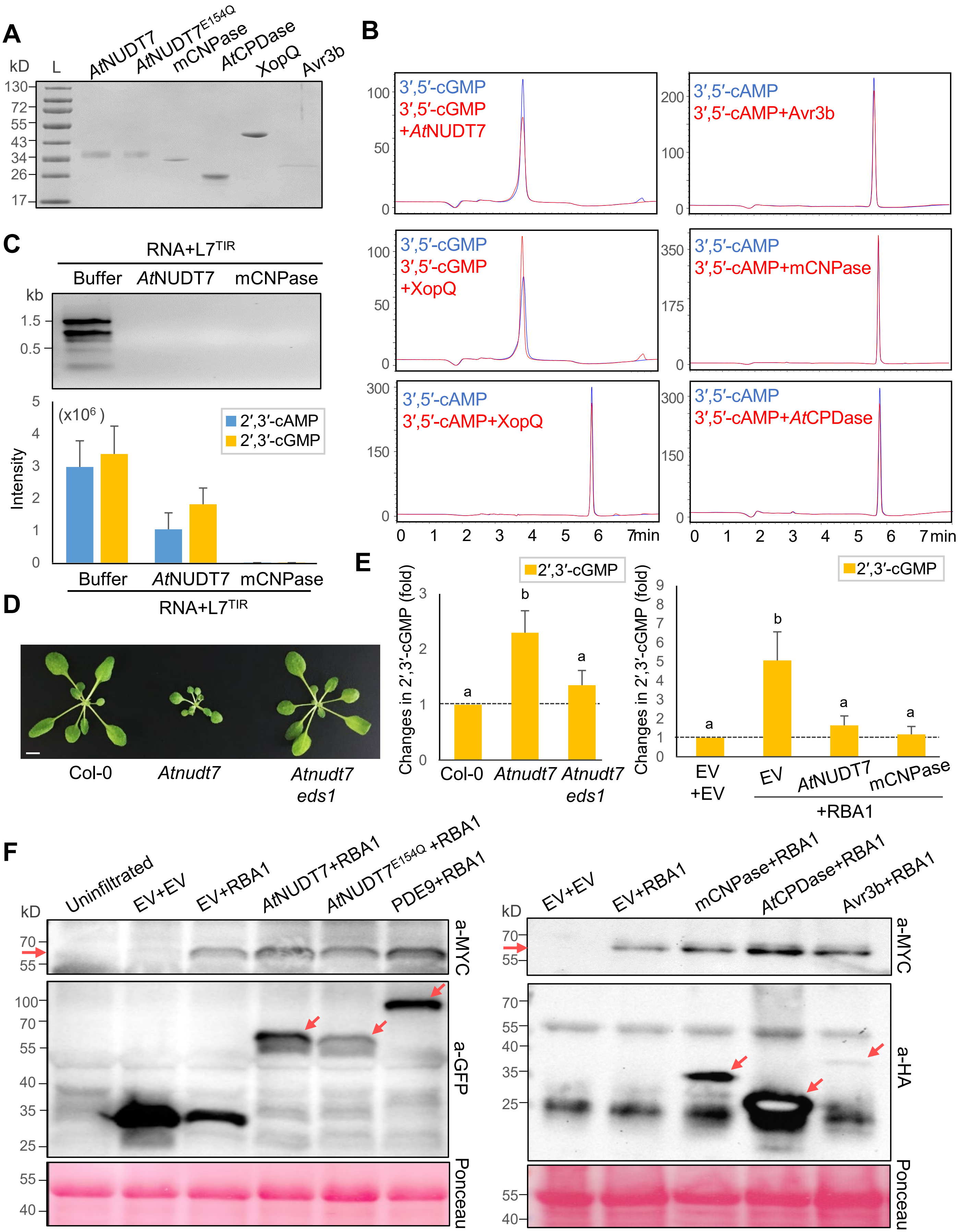

A
kD L AtNUDT7 AtNUDT7E154Q mCNPase AtCPDase XopQ Avr3b
130 72 55 43 34 26 17
B
3′,5′-cGMP
3′,5′-cGMP +AtNUDT7
3′,5′-cAMP
3′,5′-cAMP+Avr3b
3′,5′-cGMP
3′,5′-cGMP +XopQ
3′,5′-cAMP
3′,5′-cAMP+mCNPase
3′,5′-cAMP
3′,5′-cAMP+XopQ
3′,5′-cAMP
3′,5′-cAMP+AtCPDase
0 1 2 3 4 5 6 7min
C
RNA+L7TIR
Buffer AtNUDT7 mCNPase
kb
1.5
0.5
(x106 )
Intensity
2′,3′-cAMP
2′,3′-cGMP
D
Col-0
Atnudt7
Atnudt7 eds1
E
2′,3′-cGMP
Changes in 2′,3′-cGMP (fold)
EV +EV
EV
AtNUDT7
mCNPase
+RBA1
F
Uninfiltrated
EV+EV
EV+RBA1
AtNUDT7+RBA1
AtNUDT7E154Q +RBA1
PDE9+RBA1
a-MYC
a-GFP
Ponceau
mCNPase+RBA1
AtCPDase+RBA1
Avr3b+RBA1
a-HA

### Figure s7

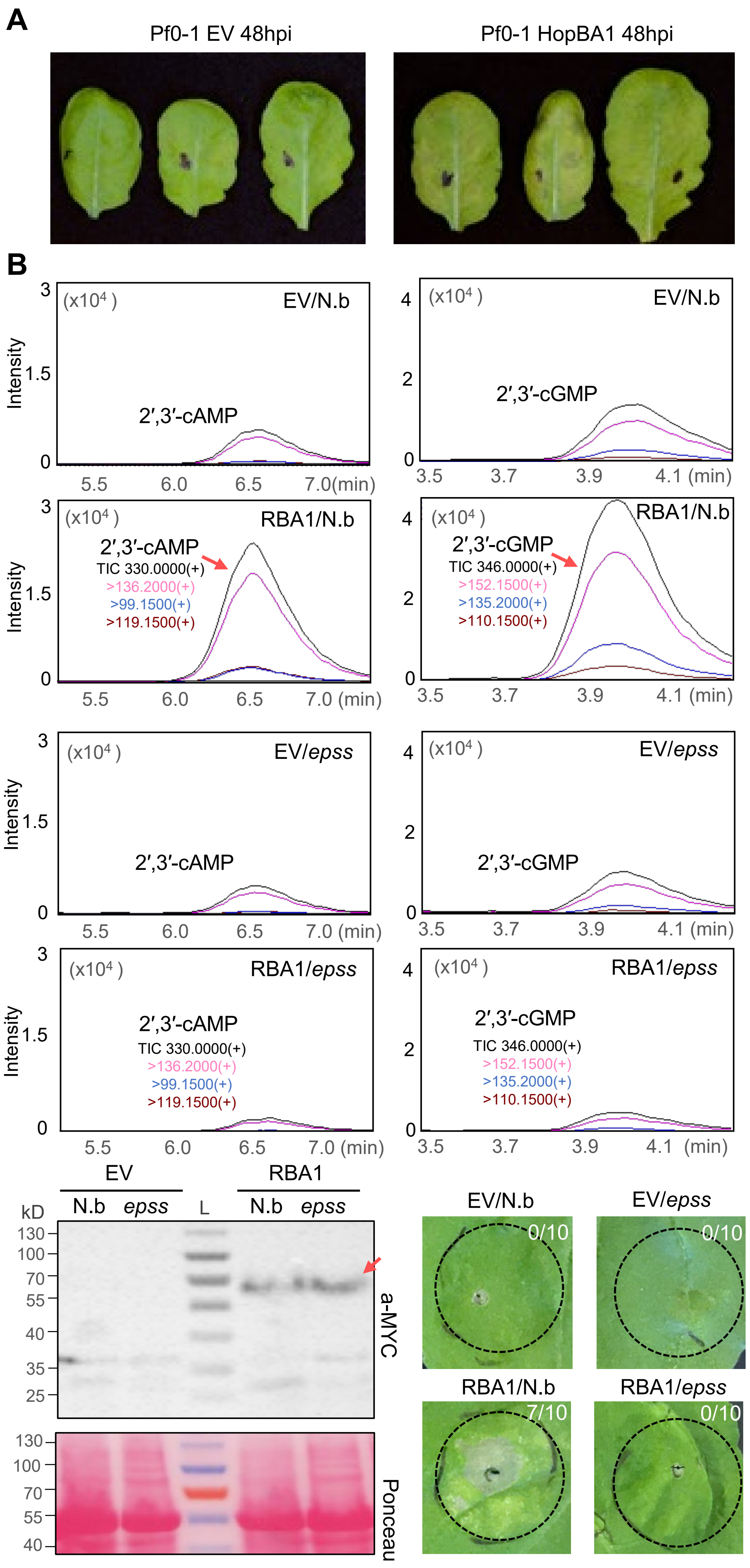

A
Pf0-1 EV 48hpi
Pf0-1 HopBA1 48hpi
B
Intensity
(x10⁴)
EV/N.b
2′,3′-cAMP
2′,3′-cGMP
RBA1/N.b
TIC 330.0000(+)
>136.2000(+)
>99.1500(+)
>119.1500(+)
TIC 346.0000(+)
>152.1500(+)
>135.2000(+)
>110.1500(+)
EV/epss
RBA1/epss
5.5 6.0 6.5 7.0(min)
3.5 3.7 3.9 4.1 (min)
EV
RBA1
N.b epss L N.b epss
kD
130 100 70 55 40 35 25
a-MYC
Ponceau
EV/N.b 0/10
EV/epss 0/10
RBA1/N.b 7/10
RBA1/epss 0/10

### Figure s7 (continued)

**C**

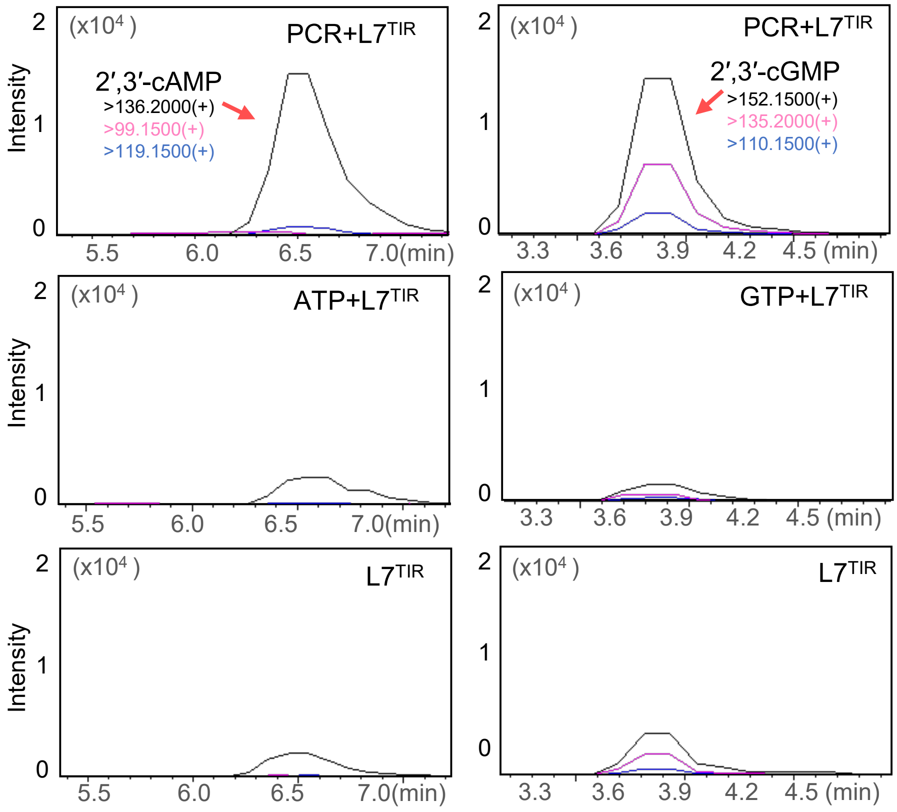


**D**

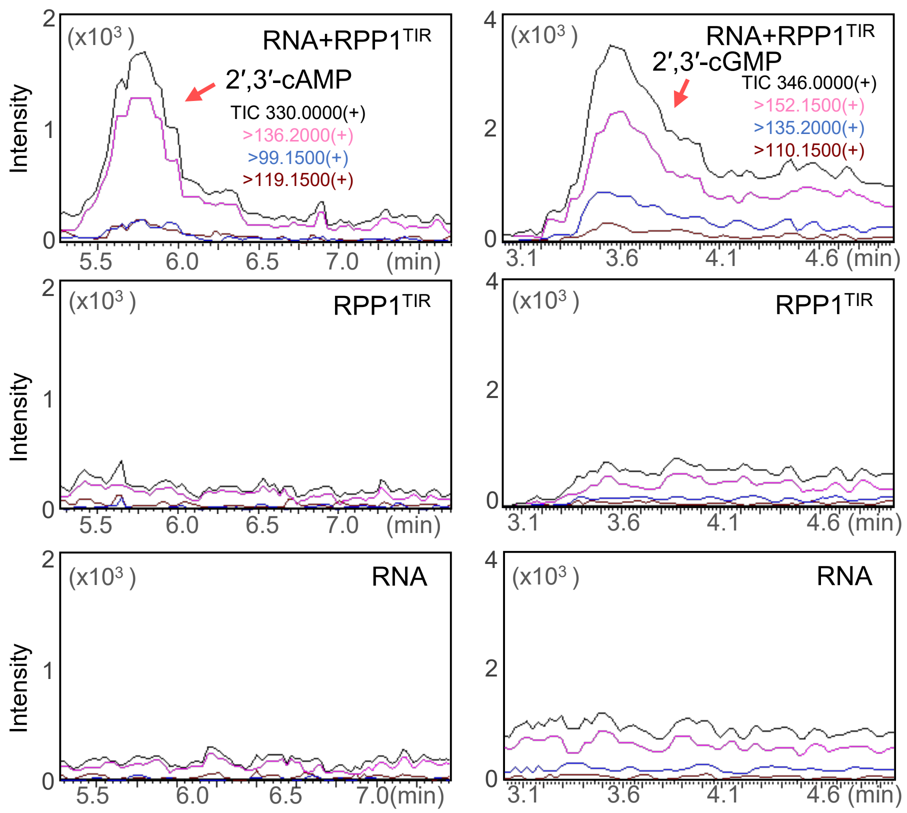
